# Efficient combinatorial targeting of RNA transcripts in single cells with Cas13 RNA Perturb-seq

**DOI:** 10.1101/2022.02.02.478894

**Authors:** Hans-Hermann Wessels, Alejandro Méndez-Mancilla, Efthymia Papalexi, William M Mauck, Lu Lu, John A. Morris, Eleni Mimitou, Peter Smibert, Neville E. Sanjana, Rahul Satija

**Affiliations:** New York Genome Center, New York, NY, USA; Center for Genomics and Systems Biology, Department of Biology, New York University, New York, NY, USA; Technology Innovation Lab, New York Genome Center, New York, NY, USA

**Author notes:** These authors contributed equally (Hans-Hermann Wessels, Alejandro Méndez-Mancilla). These authors share correspondence.

## Abstract

Pooled CRISPR screens coupled with single-cell RNA-sequencing have enabled systematic interrogation of gene function and regulatory networks. Here, we introduce Cas13 RNA Perturb-seq (CaRPool-seq) which leverages the RNA-targeting CRISPR/Cas13d system and enables efficient combinatorial perturbations alongside multimodal single-cell profiling. CaRPool-seq encodes multiple perturbations on a cleavable array which is associated with a detectable barcode sequence, allowing for the simultaneous targeting of multiple genes. We compared CaRPool-seq to existing Cas9-based methods, highlighting its unique strength to efficiently profile combinatorially perturbed cells. Finally, we apply CaRPool-seq to perform multiplexed combinatorial perturbations of myeloid differentiation regulators in an acute myeloid leukemia (AML) model system and identify extensive interactions between different chromatin regulators that can enhance or suppress AML differentiation phenotypes.

---

Recent technological advances that couple pooled genetic perturbations with scRNA-seq or multimodal characterization (i.e. Perturb-Seq, CROP-seq, CRISP-seq, and ECCITE-seq ^1–4^), promise to transform our understanding of gene function. In particular, the ability to perform combinatorial perturbations represents an opportunity to decode complex regulatory networks, with pioneering work demonstrating the ability to identify epistasis and other genetic interactions ^5–7^. However, there are specific technical and analytical challenges associated with pooled single-cell screens which are exacerbated when considering combinatorial perturbations. For example, undetected or incorrectly assigned sgRNAs can affect up to 20% of cells ^6^, but this is compounded when multiple independent sgRNAs are introduced and independently detected in each cell. Moreover, perturbations introduced by Cas9 are not uniformly efficient, and a considerable fraction of targeted cells may exhibit no phenotypic effects of perturbation ^8,9^. Therefore, when performing two or more simultaneous perturbations, the fraction of cells where all perturbations are both successfully introduced and successfully detected can decrease dramatically.

Type VI CRISPR Cas proteins, such as the VI-D family member *Rfx*Cas13d, are programmable RNA-guided and RNA-targeting nucleases that enable targeted RNA knockdown. Notably, *Rfx*Cas13d is also capable of processing a CRISPR array into multiple mature CRISPR RNAs (crRNAs) ^10^, presenting an attractive option for combinatorial perturbations at the RNA level. Recently, we confirmed that *Rfx*Cas13d can lead to striking target-RNA knockdown, and learned a set of optimal targeting rules from thousands of gRNAs tiling different transcripts ^11^. We therefore sought to combine pooled CRISPR-Cas13 screens with single-cell readouts to perform combinatorial and multimodal pooled genetic screens.

Our method for Cas13 RNA Perturb-Seq (CaRPool-seq) is enabled via an optimized molecular strategy to deliver individual or multiple gRNA perturbations in each cell and detect their identity during a single-cell sequencing experiment. Type VI A, C, and D Cas13 crRNAs consist of a short 5’ direct repeat (DR) and a variable spacer (also called guide RNA; gRNA) at the 3’end, and therefore lack a common priming site for reverse transcription. We developed an approach for direct gRNA detection by adding a 10x Genomics-compatible ‘capture’ sequence on the 3’ end of the 23nt target spacer (**Figure 1a**). We also explored an ‘indirect’ capture strategy, where a dedicated crRNA of the CRISPR array contains an array specific barcode (barcode gRNA; bcgRNA), and tested different positional configurations of the bcgRNA and gRNA (**Figure 1a**). We evaluated the performance of each method by targeting cell surface proteins and measuring knockdown via flow cytometry (**Figure 1b** and **Supplementary Figure 1**), and by quantifying crRNA detection via RT-PCR (**Figure 1c**). While all methods successfully induced robust knockdown (**Figure 1b**), we found that indirect guide capture with an optimized configuration resulted in the strongest crRNA transcript detection ability (Configuration X; **Figure 1c**).

**Figure 1:**
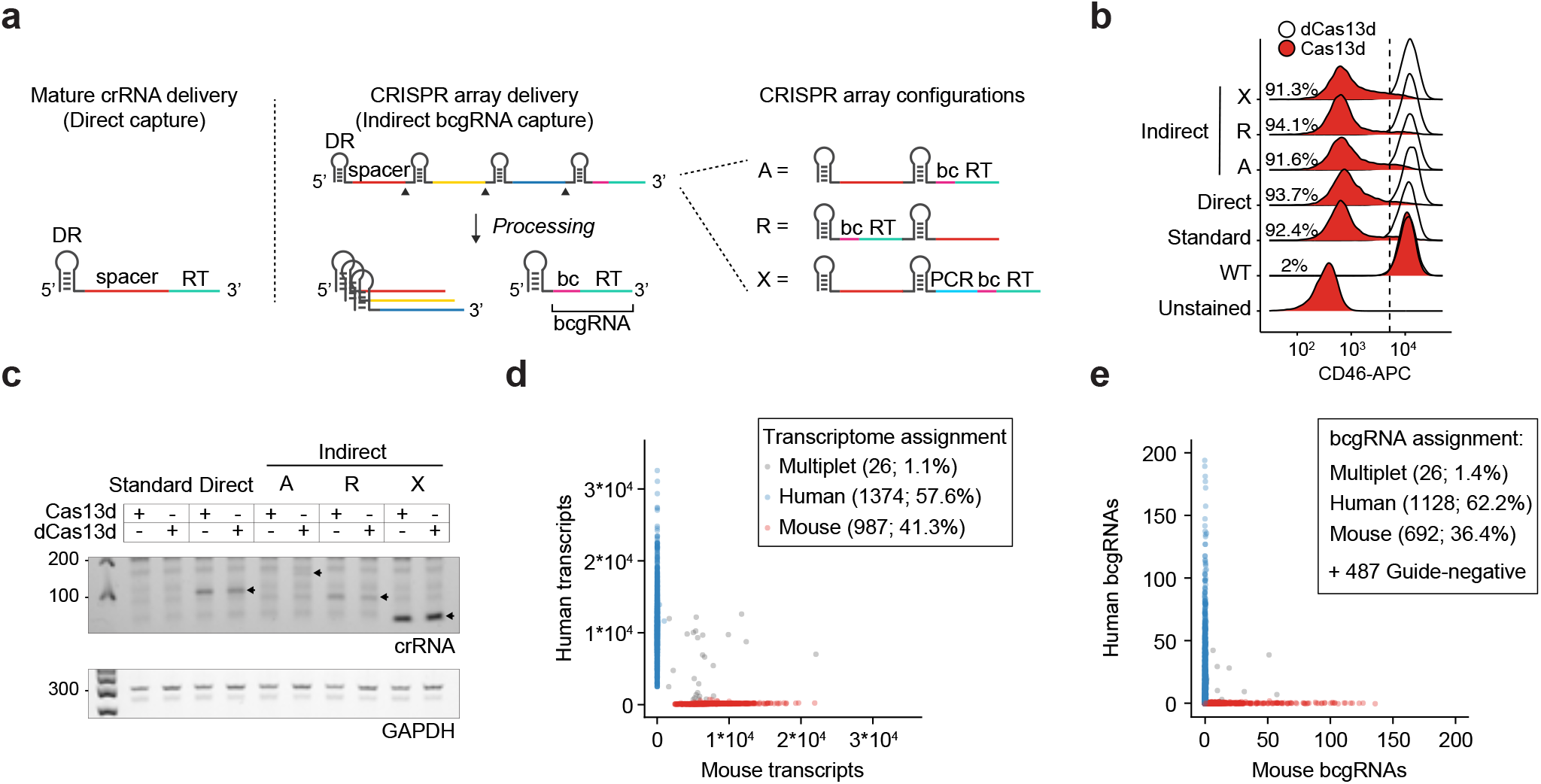
Efficient capture of gRNAs for Cas13 RNA Perturb-seq (CaRPool-seq) a) Scheme of direct and indirect array-based gRNA capture approaches. Four approaches have been assessed. Direct capture uses a reverse transcription (RT) handle added directly downstream to the spacer RNA. For the indirect capture method, a barcode guide RNA (bcgRNA) is captured as part of a CRISPR array. Three different CRISPR array configurations (A, R and X) have been tested (RT = Reverse transcription handle, bc = Barcode, PCR = PCR primer annealing site, bcgRNA = barcode guide RNA, A = Array, R = Reversed Array configuration, X = Extra PCR handle). b) Density plots showing the CD46-APC signal upon Cas13d-mediated CD46 knockdown (red) and dCas13d-mediated controls (white) using the four CaRPoool-seq configurations described in (A), as well as standard gRNA. The CS1 RT handle was used in all cases. c) PCR amplicons of reverse-transcribed crRNAs from lentivirally infected cells used in b. Direct capture (expected 109 bp), R-type array (expected 99 bp), and X-type array (expected 52 bp) allowed for reverse transcription and amplification at varying efficiency. X-type arrays are independent of template switching success and show the highest detection sensitivity. Type-X band intensities were equal between active RfxCas13 and inactive RfxdCas13d suggesting that Cas13 RNA targeting activity does not reduce available bcgRNA amounts. A-type arrays showed a faint band at the length of unprocessed CRISPR arrays (159 bp). d) Species mixing experiment profiling 2,387 HEK293FT-Cas13d or mouse NIH/3T3-Cas13d cells lentivirally transduced with CRISPR array virus. The CRISPR array includes a non-targeting gRNA and a bcgRNA in X-type configuration. The plot shows the number of transcripts associated with each cell barcode. Datapoint colors and boxed labels are assigned based on transcriptome classification (>90% species-specificity required for assignment). eNumber of bcgRNAs associated with each cell barcode. Datapoint colors are based on transcriptome classification, and boxed labels are based on observed gRNA (>90% species-specificity required for assignment).

These results demonstrate that *Rfx*Cas13d crRNAs can be modified by adding a common RT handle either directly to the gRNA or as a separate bcgRNA as part of a CRISPR array, allowing for reverse transcription and amplification. Notably, our strategy for indirect detection is well-suited for delivering multiple gRNAs into a single cell alongside a detectable bcgRNA that encodes the collective identity of these perturbations. In addition, utilizing a unique set of reverse transcription handle and Illumina PCR priming sequence in our modified crRNA (**Supplementary Figure 2**) ensures that these perturbations can be detected not only alongside scRNA-seq, but also when profiling additional molecular modalities (e.g. CITE-seq ^12^ for simultaneous transcriptome and surface protein profiling).

As proof of principle, we first tested the ability of CaRPool-seq to detect and assign bcgRNAs in a single-cell species mixing experiment. We separately transduced *Rfx*Cas13d-expressing human HEK293FT and mouse NIH/3T3 cells with a viral pool of three CRISPR arrays containing a non-targeting (NT) gRNA and a species-specific bcgRNA. We profiled a mixture of human and mouse cells with the 10x Genomics Chromium system (v3), aiming to detect both cellular transcriptomes and the bcgRNAs. Of 2387 cells, we found that 78.5% expressed a single bcgRNA (1.1% >1 bcgRNAs; 20.4% no detected bcgRNA). Moreover, we observed extremely high concordance between RNA and bcgRNA labels in singlet cells (99.2%) (**Figure 1d-e**). These numbers demonstrate that CaRPool-seq enables pooled perturbation screens that can be efficiently and accurately demultiplexed into a single-cell readout.

Next, we tested the ability of CaRPool-seq to distinguish combinatorial perturbations on multiple molecular modalities at single-cell resolution. We designed gRNAs targeting three cell surface proteins, CD46, CD55, and CD71, as well as NT gRNAs. We created 29 crRNA arrays (**Supplementary Table 1**), each of which contains up to three gRNAs and a bcgRNA, allowing for the perturbation of these genes individually or in combination. We transduced HEK293FT cells with a viral pool of all crRNAs and performed CaRPool-seq with CITE-seq ^12^ readout (**Figure 2a** and **Supplementary Figure 3a**), allowing the assessment of each perturbation on both the cellular transcriptome and antibody-derived tags (ADTs) associated with CD46, CD55, and CD71 surface protein levels.

**Figure 2:**
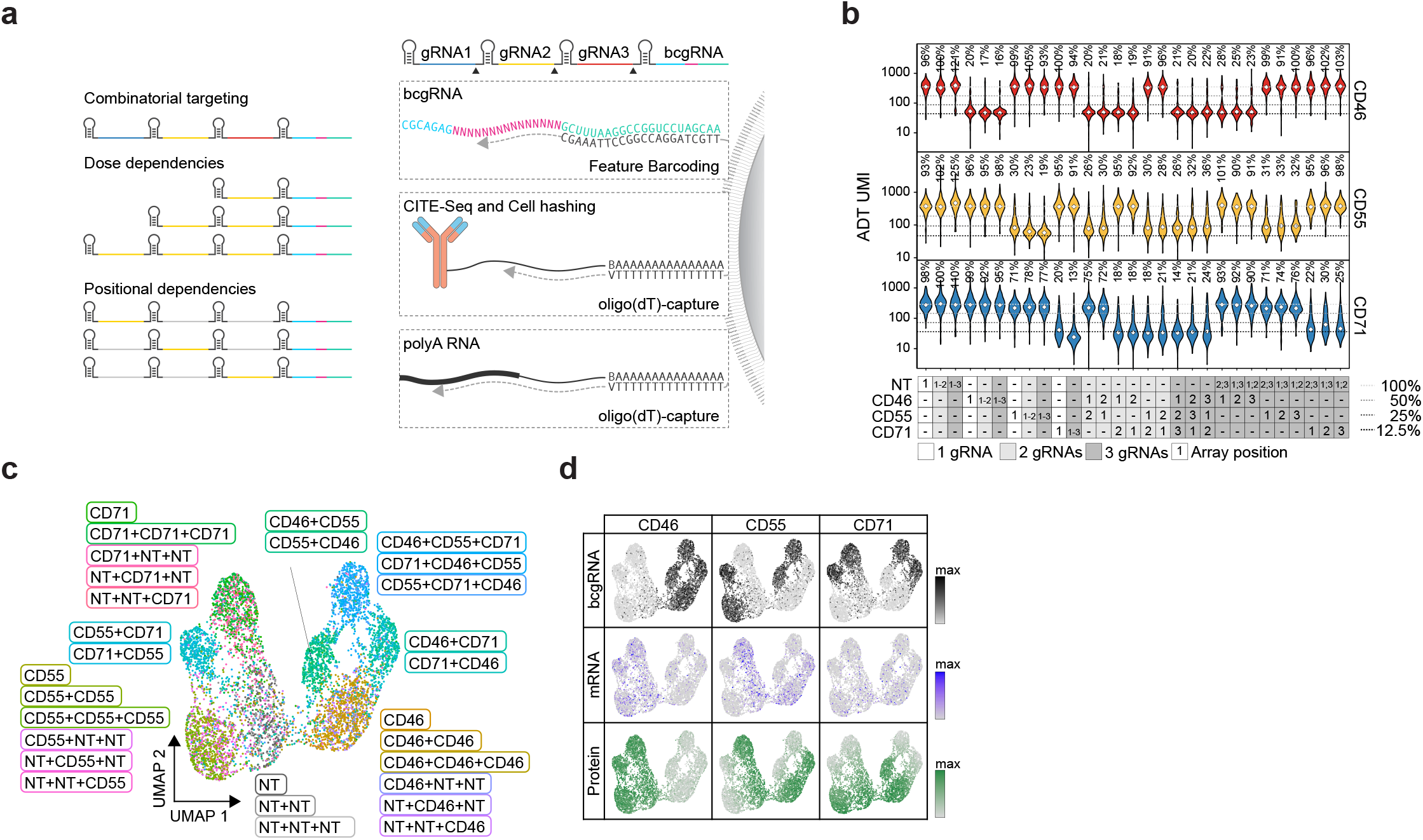
CaRPool-seq enables combinatorial gene targeting with a multimodal single-cell readout. a) CaRPool-seq can be combined with CITE-seq and Cell Hashing modalities. b) Violin plots depicting protein expression of target genes (ADT UMI counts for CD46, CD55, CD71), grouped by CRISPR arrays (n=29). Three dashed lines indicate 50%, 25%, and 12.5% UMI count relative to the median of all non-targeting cells. Diamonds indicate the median UMI count. The number above each violin plot indicates the mean level of reduction across single cells. CD71+CD71 was not included in the experiment. c) UMAP visualization of single-cell protein expression profiles of CaRPool-seq experiment (n = 6,986 cells). Cells are colored based on the single or combinatorial perturbations they received. d) Expression levels of bcgRNA (black), mRNA (blue), and protein (ADT; green) for CD46, CD55, CD71 superimposed on the UMAP visualization (n = 6,986 total cells).

We obtained 9,355 single-cell profiles and demultiplexed them into groups based on the detected bcgRNA (**Supplementary Figure 3b**; 74.7% expressed a single bcgRNA, 80.8% expressed at least one bcgRNA). We observed, on average, a 76.5% (+/− 5.7%) mean reduction in protein levels for each targeted gene after perturbation with Cas13 demonstrating clear evidence of robust molecular perturbation (**Figure 2b-d**). Moreover, the strength of knockdown was similar for multi-gRNA crRNA arrays relative to single gRNA perturbations (**Figure 2b** and **Supplementary Figure 4**). When examining transcriptomic pseudobulk profiles for all 26 targeting gRNA groups, we observed decreased mRNA expression for each targeted transcript, even when perturbing transcripts of three genes simultaneously (15 examples in **Supplementary Figure 3c-d**). The average strength of transcriptomic knockdown (mean 15%, sd 5.2%) was consistently reduced compared to the observed protein reduction. This is concordant with the mode of knockdown exhibited by Cas13. Target RNAs are continuously being produced and degraded before the target can be translated into protein. Further, analogous to how Cas9-nuclease targeting often produces RNAs degraded by nonsense mediated decay ^15^, it is possible that Cas13 cleavage produces RNA molecules that can be detected by scRNA-seq but cannot be translated into functional protein, suggesting that the level of measured RNA knockdown underestimates the phenotypic effect of Cas13 perturbation. Importantly, we found that Cas13d-mediated gene knockdown was highly specific, with no evidence of off-target effects in any of the three target gene perturbations (**Supplementary Figure 3e-f**).

We next benchmarked the performance of CaRPool-seq against direct capture Perturb-Seq ^6^ using three different Cas9 effectors: Cas9-nuclease, a first-generation CRISPR inhibition (CRISPRi) system, KRAB-dCas9 ^16,17^ and a second-generation, dual-effector CRISPRi system, KRAB-dCas9-MeCP2 ^17,18^. In CaRPool-seq one bcgRNA encodes the combined gRNA identities, while direct capture Perturb-Seq requires independent detection of one sgRNA feature per perturbation (**Figure 3a**). We replicated our previously described experimental system, targeting the same three cell surface markers (CD46, CD55, and CD71) alone or in combination. For each target, we evaluated three sgRNAs from established CRISPR-KO ^19^ and CRISPRi ^20^ sgRNA libraries (**Supplementary Figure 5a**) and selected the best sgRNA for Perturb-Seq (**Supplementary Figure 5b**). In addition, we utilized Cell Hashing ^21^ to label cells targeted with vectors encoding single, double, or triple perturbations. As in CaRPool-seq, we quantified gRNA, RNA, and ADT levels in each cell.

**Figure 3:**
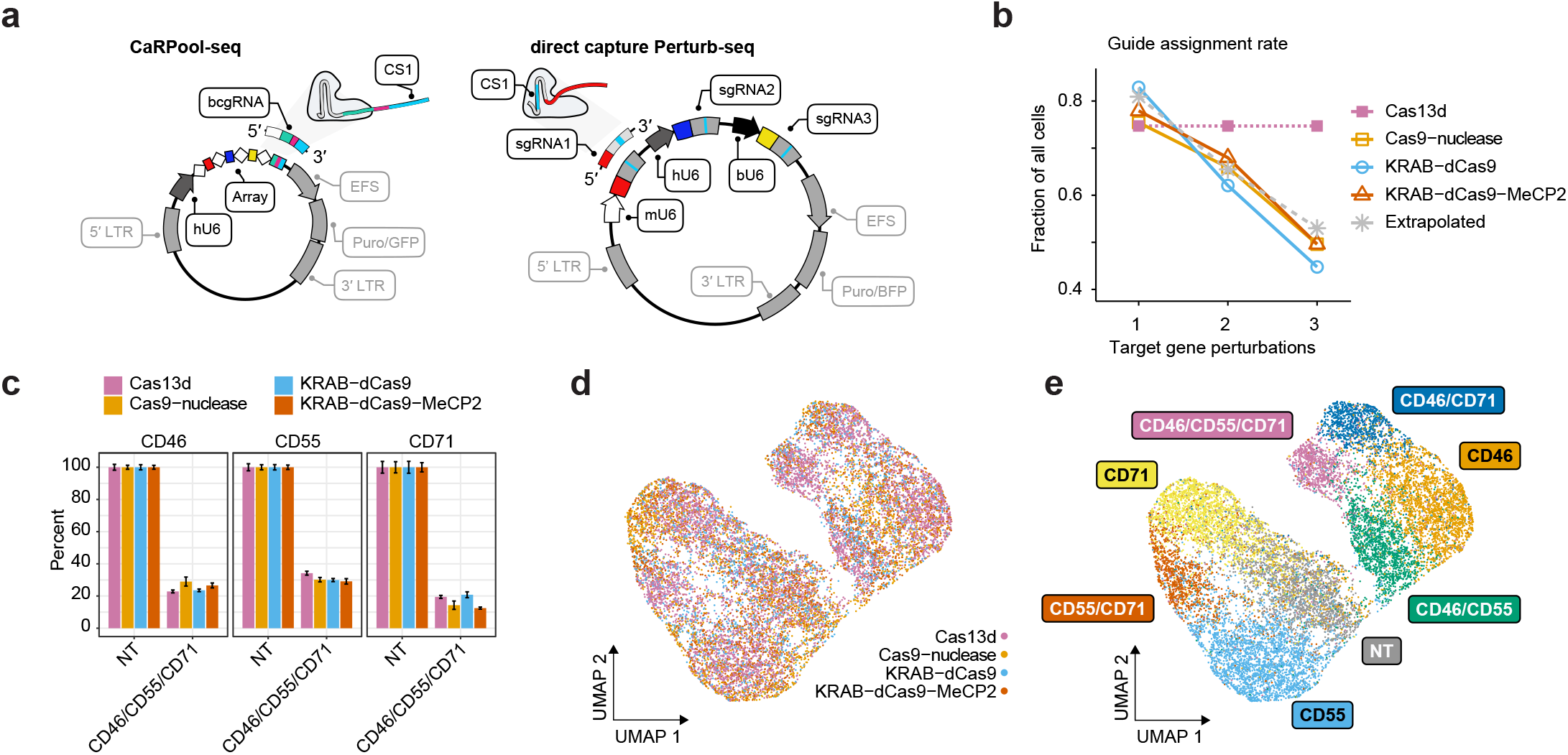
Benchmarking CarPool-seq against alternative combinatorial perturbation approaches. a) Plasmid vectors for lentivirus production for triple perturbation scenarios comparing CaRPool-seq and Direct Capture Perturb-seq. In both cases, the feature of interest is captured using 10x Genomics feature barcoding technology. In CaRPool-seq a single barcode sequence (bcgRNA) represents a combinatorial perturbation, while Perturb-seq requires the independent capture of multiple gRNA. b) Fraction of cells where the correct combination of gRNA was detected for single, double, and triple perturbations. The dashed grey line represents a theoretical extrapolation based on an assumption of independent sgRNA detection with p=0.81, as reported. in Replogle et al. 6. c) Relative expression of cell surface proteins CD46, CD55, and CD71 in cells with assigned non-targeting (s)gRNAs (NT) or a combination of three targeting (s)gRNAs. The expression level of each target is normalized to NT control. Bars indicate mean across cells with s.e.m. error bars. d) Protein level ADT-based clustering of single-cell expression profiles of merged CaRPool-seq, Perturb-seq experiments using Cas9, KRAB-dCas9, or KRAB-dCas9-MeCP2. Cells are colored by perturbation technology, and in e) cells are colored based on the single or combinatorial perturbation received. Cells cluster together across technologies, indicating that all approaches introduce perturbations of similar phenotypic strength.

Our benchmarking analysis found that, in contrast to CaRPool-seq, alternative Cas9-based approaches struggled to efficiently identify and detect combinatorial perturbations (**Figure 3b**). For example, in the KRAB-dCas9-MeCP2 experiment, we recovered 1,570 cells that received vectors targeting three genes. Among these cells, only 779 (49.6%) were associated with the correct three sgRNA after sequencing. In the remaining cells, we detected too few perturbations (0, 1, or 2 gRNAs, 31.2%), too many (4+ gRNAs 10.0%), or an improper combination of 3 gRNA (9.2%). This observed drop-off is fully consistent with the theoretical expectation of recovery for multiple independently detected gRNAs and highlights the challenge of efficiently profiling multiple perturbations with existing approaches. Since CaRPool-seq associates combinatorial perturbations with a single bcgRNA, the efficiency of detection does not vary between single and multiple perturbations.

We next compared the strength of perturbation across methods. We first considered cells where three perturbations were successfully detected based on either the bcgRNA (CaRPool-seq) or independently detected gRNA (Perturb-Seq). When considering these cells, all methods successfully induced a similarly strong depletion of all three surface proteins (Cas13d: 74.5%, Cas9; 75.5%, KRAB-dCas9; 75.2%, KRAB-dCas9-MeCP2; 77.3%) (**Figure 3c**). We next analyzed all cells based on their ADT levels. CaRPool-seq and Perturb-Seq cells clustered together (**Figure 3d**), and grouped by gRNA identity (**Figure 3e** and **Supplementary Figure 5c**), again demonstrating that the strength of phenotypic protein perturbation was similar across all methods. We conclude that CaRPool-seq and Perturb-Seq can both effectively introduce combinatorial perturbations into single cells. However, CaRPool-seq exhibits clear advantages in the ability to successfully identify and detect these perturbations and therefore represents an attractive approach for performing combinatorial single-cell CRISPR screens.

To demonstrate the throughput and potential of CaRPool-seq to characterize genetic interactions, we performed a multiplexed screen of 158 combinatorial gene pairs. Motivated by recent work ^22^, we aimed to characterize potential interactions between previously identified regulators of leukemic differentiation, which can influence the response to chemotherapy and small-molecule drugs. We generated a human MLL-AF9 NRAS^G12D^ AML cell line (THP-1 cells), with a stably integrated doxycycline-inducible Cas13d cassette, as a model system. We first performed a bulk Cas13d CRISPR screen using a targeted library of 439 genes with 10 gRNA per gene. On day 13-16 post Cas13d-induction, cells were sorted into bins based on their surface expression of CD14 and CD11b, immunophenotypic markers of monocyte differentiation. By comparing gRNA representation between low and high-expressing bins, we selected 26 genes (and for each, the two best performing, non-overlapping gRNAs) that influenced differentiation (**Supplementary Figure 6a-i**). Through individual perturbations with a flow-cytometry readout, we validated that each of these genes regulated CD11b expression (**Supplementary Figure 6j**). Consistent with previous work ^22^, these genes were largely associated with DNA-binding and chromatin remodeling functions, and include a subset of previously identified regulators of AML differentiation.

We next applied CaRPool-seq to test the effects of combinatorially perturbing these regulators (**Figure 4a, Supplementary Figure 7a-b**). We infected cells with a pooled library of 385 crRNA arrays. This library encoded 28 single perturbations (26 regulators and two negative control genes) and 158 paired perturbations. It also encompassed technical replicates for each perturbation using independent gRNAs, as well as NT controls. We profiled the transcriptome, cell surface protein levels, and gRNA expression for 31,308 demultiplexed single cells.

**Figure 4:**
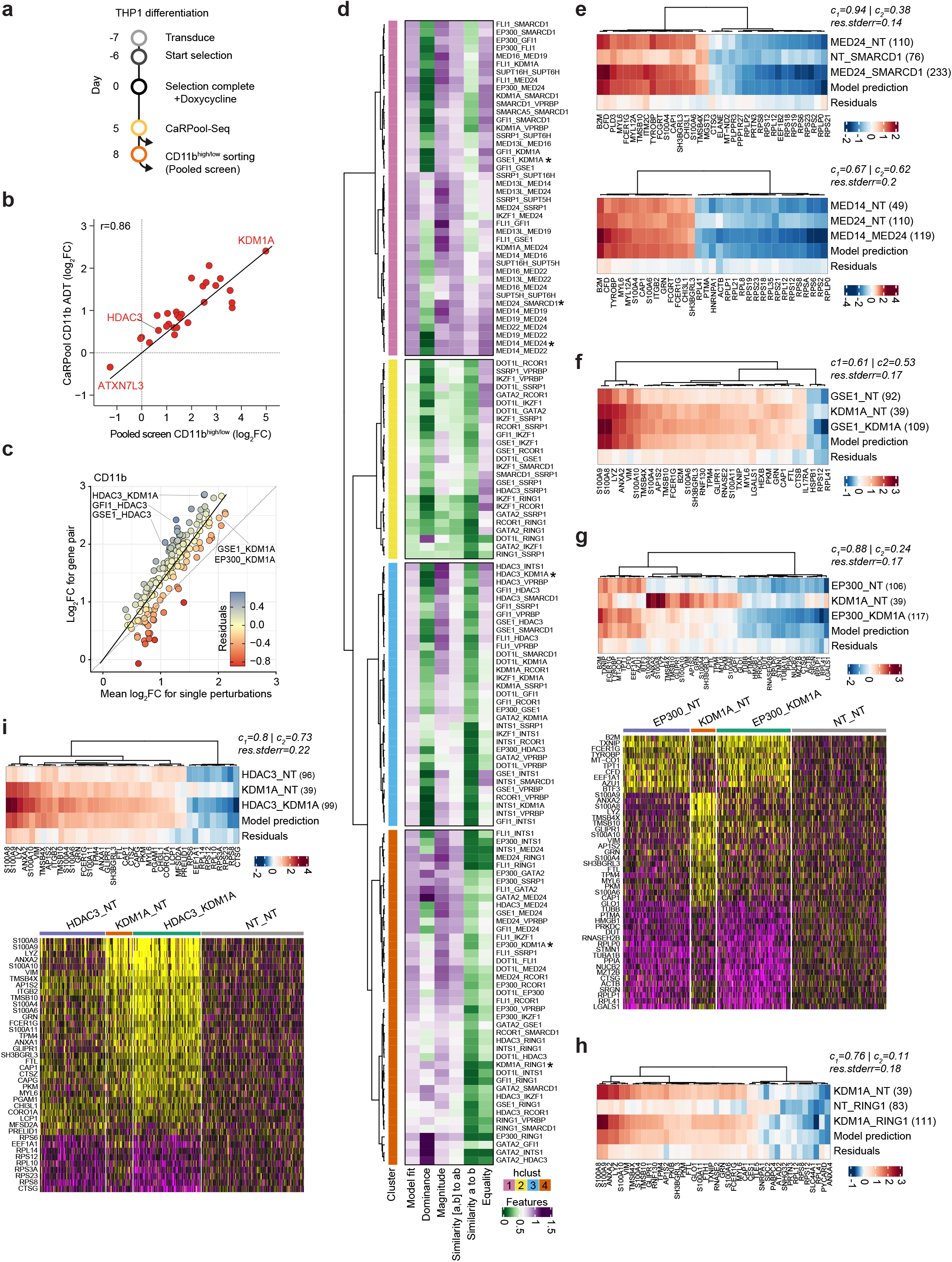
CaRPool-seq identified genetic interactions in AML differentiation. a) Timeline for THP1 cell infection and CaRPool-seq and pooled screen readout for CD11b fluorescent activated cell sorting (FACS). We transduced a pooled lentivirus library with 385 bcgRNA CRISPR arrays, containing 158 gene pair perturbations, 28 single gene perturbations (each with two indepen-dent technical replicates), and 13 non-targeting controls. b) Correlation of bcgRNA enrichment in the pooled screen (CD11bhigh/low) and CD11b ADT log2FC for cells grouped by assigned bcgRNA cells relative to non-targeting control cells in CaRPool-seq. c) Correlation of CD11b ADT log2FC of in cells with double perturbations, and the mean log2FC of both single perturbations (n=158 gene pairs). Residuals indicate the distance to the average linear relationship. d) Regression model results decomposing the observed perturbation responses in doubly perturbed cells as a linear combination of single gene perturbation responses. Dual perturbation responses are clustered by their scaled model fit coefficients (see methods). e-i) Comparison of transcriptional responses for double vs single perturbation. Heatmaps show deviation in average gene expression relative to unperturbed cells for the 20 most significantly regulated genes (Wilcoxon test). These heatmaps visualize a range of observed interactions between gene pairs, including cases where genes contribute equally to the dual perturbation response (e, f), and where one gene’s perturbation signature dominates over the other (g-i). Average heatmaps in g and i are accompanied by single-cell gene expression heatmaps.

We first compared the level of surface protein expression for each perturbation to NT controls (**Supplementary Figure 7c**). As expected, we found that each single-gene perturbation affected CD11b expression, with observed log_2_-fold changes that were in strong agreement (R=0.86; **Figure 4b** and **Supplementary Figure 7d-f**) with the level of gRNA enrichment from bulk CRISPR screens. Observed log_2_-fold changes for all perturbations were also reproducible (R=0.82; **Supplementary Figure 7g**) across technical replicate perturbations. We next compared the observed effect of the 158 dual gene perturbations to the effects resulting from the two single perturbations. We observed a strong correlation and found that the dual perturbation was typically stronger than the average of individual knockdowns, but weaker than the product. We also observed both synergistic and dampening effects. For example, individual knockdown of the histone demethylase KDM1A (log_2_FC 2.41) and the histone deacetylase HDAC3 (log_2_FC 0.53) lead to strong and weak CD11b-upregulation, but dual perturbation leads to a synergistic effect (log_2_FC 2.85). In contrast, while individual knockdown of EP300 also leads to CD11b upregulation (log_2_FC 1.57), dual perturbation with KDM1A (log_2_FC 2.05) was weaker than the individual KDM1A knockdown. We validated these findings using data from our Cas13d CD11b pooled screen (**Supplementary Figure 7 d-f** and **h**).

We next explored the transcriptional profiles of both single and double perturbations in our CaRPool-seq dataset. As expected, up-regulated gene modules in singly perturbed cells were typically associated with genetic programs associated with the differentiation and function of myeloid cells (**Supplementary Figure 7i**). We applied a recent pioneering framework ^7^ that fits a regression model to decompose the observed perturbation responses in doubly perturbed cells as a linear combination of single gene perturbation responses. The fit and coefficients of this model describe multiple types of genetic interactions, including epistasis, genetic suppression, and synergistic relationships. Fitting these models to each of our pairwise perturbations revealed a diversity of genetic interactions, which we broadly clustered into 4 groups (**Figure 4d**). For 33 gene pairs in Cluster 1, we saw that each individual gene’s profile contributed equally to the dual perturbation response, and the linear model exhibited a strong fit. As a positive, control, many of the pairs in this cluster represented perturbations of two proteins in the same complex (i.e. MED14/MED24; SUPT16H/SUPT6H) (**Figure 4d-e**). This cluster also represented pairs of proteins residing in separate complexes (MED24/SMARCD1 of mediator and SWI/SNF complexes) which share similar perturbation signatures (**Figure 4e**). Dual perturbation of KDM1A and the transcriptional repressor GSE1 also fell in this cluster (**Figure 4f**), consistent with previous work that suggests a cooperative interaction via colocalization at repressed promoters to inhibit myeloid differentiation ^23^.

In cluster 4, we identified genetic interactions where one gene’s effect appeared to dominate over the other. We generally observed that transcriptional responses varied widely when pairing KDM1A knockdown with different chromatin regulators. For example, we found that the EP300-signature appeared more strongly than the KDM1A-signature when combinatorial perturbing both genes (**Figure 4g**). Dually perturbed cells exhibited higher expression of progenitor genes (i.e. the progenitor marker AZU1), and reduced expression of differentiated marker genes (myeloid marker S100A4) compared to individual KDM1A perturbation. Contrastingly, the KDM1A response signature dominated when paired with perturbation of the polycomb repressive complex member RING1 (**Figure 4h**). Dual perturbation of HDAC3 enhanced the KDM1A transcriptional response signature, consistent with our previously described immunophenotypic results for these cells (**Figure 4i**). These findings also support and provide a molecular explanation for recent observations that combination therapies of KDM1A antagonist and HDAC inhibitors exhibit an enhanced response ^24^. The heterogeneity across interaction responses was not unique to KDM1A, but describes many genetic regulators in our study, and highlights CaRPool-seq’s ability to robustly characterize complex genetic interactions at scale.

## Discussion

Here, we present CaRPool-seq, a flexible method for performing CRISPR Cas13 RNA-targeting screens with a single-cell sequencing-based readout. We introduced an optimized strategy to deliver multiple gRNA as part of a single CRISPR array, which is subsequently cleaved into individual crRNAs. We demonstrate that this strategy is well-suited for performing combinatorial perturbations, whose identity is encoded in a single barcode that can be reliably detected alongside multiple molecule modalities including scRNA-seq and CITE-seq.

Through benchmarking, we show that CaRPool-seq is more efficient and accurate when assigning multiple perturbations in single cells when compared to Cas9-based technologies. Even with individual perturbations, user will still benefit from CaRPool-seq. In particular, as an RNA-targeting enzyme, Cas13d can be uniquely applied to target specific RNA isoforms, or even circular, enhancer, or antisense RNA molecules. RNA-directed approaches may also be optimal when targeting a single member of a local gene cluster, where alternative KRAB-mediated repressive strategies may ‘spread’, and introduce off-target effects ^25^. CaRPool-seq can profile additional cellular modalities such as cell surface protein levels and, in the future, can be extended to additional molecular modalities including intracellular protein levels and chromatin accessibility.

Combinatorial screens have the potential to shed substantial new light on the structure of genetic regulatory networks, but also to identify combinatorial perturbations that achieve desirable cellular phenotypes. Our CaRPool-seq analysis of AML differentiation regulators benefited from recently developed computational frameworks to identify genetic interactions from multiplexed perturbation screens, and these types of data will be valuable resources for systematic reconstruction of complex pathways and cell circuits. Moreover, our identification of combinatorial perturbations that enhanced AML differentiation phenotypes was consistent with previous identification of efficacious multi-drug therapies, suggesting that future experiments may help to nominate candidates for combined drug treatments. We conclude that CaRPool-seq represents a powerful addition to the growing toolbox of methods for multiplexed single-cell perturbations.

## AUTHOR CONTRIBUTIONS

HHW, NES, and RS conceived the research. RS and NES supervised the research. HHW and AMM performed experimental work, assisted by EP, WMM, and LL. EM and PS advised with bcgRNA design. JAM validated dCas9-effector constructs. All authors participated in data interpretation. HHW, NES, and RS wrote the manuscript with input from all authors. All authors read and approved the final manuscript.

## ACKNOWLEDGEMENTS

We thank the Technology Innovation lab as well as all members of the Sanjana and Satija labs for helpful discussions. We are grateful to Zharko Daniloski for cloning the Cas9-effector protein plasmids, to Ioannis Aifantis for advice and helpful discussion related to THP1 experiments, and the Technology Innovation lab for generously sharing CITE-seq reagents. NES and RS are supported by New York University and New York Genome Center startup funds. NES is further supported by DARPA (D18AP00053), the Brain and Behavior Foundation, the Cancer Research Institute, the National Institutes of Health (NIH)/National Human Genome Research Institute (DP2HG010099), and the NIH/National Cancer Institute (R01CA218668).RS is supported by the Chan Zuckerberg Initiative (EOSS-0000000082 to RS, HCA-A-1704-01895 to PS and RS), and the National Institutes of Health (DP2HG009623-01 to RS, RM1HG011014-01 to PS and RS).

## DATA AND CODE AVAILABILITY

Raw and processed sequencing data has been submitted to Gene Expression Omnibus (GEO accession number pending).

## METHODS

### Cell culture and monoclonal cell line generation

HEK293FT cells were acquired from ThermoFisher (R70007), NIH/3T3, and THP1 cells were obtained from ATCC (CRL-1658, TIB-202). HEK293FT and NIH/3T3 cells were maintained at 37°C with 5% CO_2_ in DMEM with high glucose and stabilized L-glutamine (Caisson DML23) supplemented with 10% fetal bovine serum (Serum Plus II Sigma-Aldrich 14009C) and no antibiotics. THP1 cells were grown in RPMI supplemented with 10% FBS at 37°C with 5% CO_2_ and no antibiotics. Doxycycline-inducible *Rfx*Cas13d-NLS HEK293FT, THP1, and NIH/3T3 cells (Addgene #138149), as well as doxycycline-inducible nuclease-inactive RfxdCas13d-NLS HEK293FT, have been generated as described before ^11^. We sorted individual suspension THP1 cells using a flow cytometer (SONY SH800) to select single clonal lines. Each THP1-Cas13d clone was evaluated to provide homogenously strong CD46 knockdown using lentiviral integration of a single gRNA, puromycin selection, and flow cytometry.

Monoclonal Cas9-effector protein-expressing HEK293FT cell lines were generated using dilution plating as described above ^11^. For each Cas9-effector cell line, we evaluated multiple clones in their ability to provide homogenous and complete CD55 knockout or knockdown using lentiviral integration of a single sgRNA expressing cassette, puromycin selection, and flow cytometry. The cloning of KRAB-dCas9 and KRAB-dCas9-MeCP2 constructs have been described before ^17^. For Cas9-nuclease, we previously cloned lentiCas9-Blast (Addgene #52962) ^26^. All monoclonal CRISPR Cas effector expressing cells were maintained using 5μg/mL Blasticidin S (ThermoFisher A1113903).

### Cloning of individual Cas13 and Cas9 guide RNAs

Cas13 guide RNA cloning was done as described previously ^26^. Specifically, we cloned gRNA or barcode oligos into pLentiRNAGuide_001 (Addgene #138150). Guide RNA constructs with reverse transcription handles on the 3’ end of the spacer sequence were synthesized and cloned together. Guide RNA constructs using CRISPR arrays used in data presented in Figure 2 were cloned stepwise by introducing a guide along with a direct repeat and reconstituted BsmBI restriction sites to allow for serial cloning and extension of CRISPR arrays.

For direct capture Perturb-seq, we cloned all sgRNA-expressing constructs stepwise. First, we cloned dual sgRNA oligos into pJR85 and pJR89 as described before ^6^. In addition, we cloned single sgRNAs into custom sgRNA expressing plasmids. We used either a human U6 (hU6) promoter driving the sgRNA scaffold described in the 10x Genomics manual CG000184 Rev C (internal CS1) (pLCR36) or using the sgRNA cassette from pJR73 ^6^ driven by a bovine U6 (bU6) promoter (pLCR67). The bU6 and sgRNA cassette from pLCR67 was then subcloned into the dual sgRNA plasmid to generate triple sgRNA expressing plasmids (see also Supplementary Figure 5b). For direct capture Perturb-seq, we used pLCR36 single sgRNA constructs for pools expecting one sgRNA, pJR85/pJR89 constructs for pools expecting two sgRNAs, and pJR85/pJR89/pLCR67 constructs for pools expecting three sgRNAs. All constructs were confirmed by Sanger sequencing. All primers used for molecular cloning and guide sequences are shown in **Supplementary Table 2**. Plasmids (pLCR36 and pLCR67) will be made available on Addgene by the time of publication.

### Pooled Cas13d library design and cloning

We design two libraries for pooled cloning, one to identify genes that lead to THP1 cell differentiation (Supplementary Figure 6), and one for combinatorial targeting with CaRPool-seq (Figure 4).

First, we designed a *Rfx*Cas13d gRNAs library for single gRNA expression targeting 439 individual genes. We selected 240 target genes that led to CD11b and CD14 upregulation in a previous Cas9-based pooled screen, in addition to 199 control genes involved in TLR4 signaling. For each, we selected the transcript with the highest isoform expression (CCLE - https://sites.broadinstitute.org/ccle/datasets) and scored possible gRNA sequences using our previously described Cas13design algorithm ^11^. For each gene, we selected 10 gRNA from efficacy quartile Q4 (or Q3 if not sufficient Q4 gRNAs were found), while trying to evenly spread the gRNAs along the coding region. We only selected gRNAs without secondary target sites with 0-2 mismatches to the gRNA sequence ^27^. In total, we designed 4390 gRNAs and added 410 non-targeting control gRNAs (>3 mismatches to any hg19-annotated transcript). The gRNA library was cloned as described before ^11^. Pooled oligonucleotides were synthesized (Twist), amplified using 8x PCR reactions with 8 amplification cycles using a direct repeat specific forward primer (**Supplementary Table 2**). The amplicon was Gibson cloned into pLentiRNAGuide_001 and pLentiRNAGuide_002 (Addgene 138150, 138151). Complete library representation with minimal bias (90^th^ percentile/10^th^ percentile crRNA read ratio: 1.8 for both libraries) was verified by Illumina sequencing (MiSeq).

To design the CaRPool-seq library, we manually inspected all gRNA enrichments from the pooled screen library described above. For the 28 selected target genes, we picked the two most enriched (or depleted for CD14 and ATXN7L3) gRNAs, while avoiding overlapping gRNAs. For each gene, we paired the two gRNA with a non-targeting gRNA (n=28 single perturbations, n=58 arrays). For 17 genes, we designed all pairwise combinations (n = 132 gene pairs, n = 264 arrays). And for 9 genes we designed a subset of possible gene pairs within the same complex (n = 22 gene pairs, n = 44 arrays). We added 13 arrays non-targeting control arrays. In total, we design 385 arrays with 186 single or double perturbation constructs, each represented by two independent technical replicate gRNA combinations. We designed random 15mer sequences with hamming distance >4 to one another. We balanced the relative CRISPR array abundance by the effect on cell proliferation of the targeted genes and increased the number of array copies in the pool to minimize dropout in the CaRPool-seq experiment. The oligos for synthesis were designed in the following way:

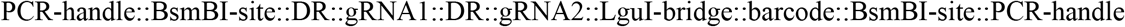

Pooled oligonucleotides were synthesized (Twist) and diluted to 1 ng/ul before amplification. The outer PCR-handles allowed for PCR amplification of the oligo pool (Supplementary Table 2) (we used Pfu-Ultra-II following the manufacturer’s recommendation using 1μl of enzyme and 20ng of oligo pool in a 50μl reaction 95C/2m, 5x[95C/20s, 58C/20s, 72C/15s], 72C/3min). The amplicon was purified using a 2x SPRI cleanup followed by BsmBI-digestion, and SPRI cleanup. All of the purified product was ligated into BsmBI-digested pLCR65 using T7-DNA ligase and cloned as described before ^11^ with > 1000 colonies per construct. The resulting plasmid pool was digested with LguI to enable ligation of the third DR and smallRNA-handle to complete the bcgRNA and CRISPR array. pLCR65 was generated by removing all LguI sites from the pLentiRNAGuide_002 plasmid, introducing the CS1 capture sequence 5’ to the terminator sequence, and replacing the puromycin gene with GFP-P2A-puromycin. The LguI insert containing the third DR and smallRNA handle was cloned into pLentiRNAGuide_001, digested with LguI, and gel-purified (2% eGel). Complete library representation with minimal bias (90^th^ percentile/10^th^ percentile crRNA read ratio: 2.6-4.8 for the relative abundance tiers and 5.0 overall) and correct gene pair to bcgRNA linkage (>94%) was verified by Illumina sequencing (MiSeq). During library cloning, we noticed two critical details: Alternative polymerase KAPA and Q5 can lead to a stronger bias in relative array abundance. Additionally, reducing the number of PCR cycles with increased oligo pool input amounts can decrease bcgRNA reassortment. Further, while we chose a two-step cloning strategy, we believe that a single step cloning strategy may yield similar results. pLCR65 will be deposited to Addgene by the time of publication.

### Virus production and viral transductions

For virus production of individual gRNA and sgRNA expressing plasmids, we seeded 1×10^6^ HEK293FT cells per well in 6-well format 12-18 hours before transfection. Per transfection reaction we used 7.5ul of 1 mg/mL polyethyleneimine (PEI) and 2.325 μg of total plasmid DNA (825 ng psPAX2: Addgene #12260; 550 ng pMD2.G: Addgene #12259; 1000 ng of gRNA/sgRNA expressing plasmid). Six to eight hours post-transfection, the medium was exchanged for 2 ml of DMEM + 10% FBS containing 1% bovine serum albumin (BSA). Viral supernatants were collected after additional 48 hours, spun down to remove cellular debris for 5 min at 4°C and 1000xg, and stored at −80C until use.

For pooled virus production the plasmid pools were transfected as described above in a 10 cm dish format (1×10^7^ HEK293FT cells, 70μl PEI, 6.4 μg psPAX2, 4.4 μg pMD2.G and 9.2 μg of the plasmid pool). The pooled virus was cleared by spinning down cell debris (3min, 1000xg) and passed through a 0.45 μm filter prior to storage at −80°C. For the CaRPool-seq experiments presented in Figures 1 and 2, and the direct capture Perturb-seq experiments presented in Figure 3, we pooled individual plasmids at equal amounts. In this way, we generated separate sgRNA pools for 1, 2, or 3 sgRNAs to a total of 6 pools (3 for Cas9-nuclease and 3 for KRAB-dCas9(-MeCP2)).

For experiments using single gRNA or CRISPR arrays, we transduced 1×10^6^ HEK293FT or THP1 cells with an MOI of 0.2-0.5. The cells were selected with 1μg/ml puromycin (ThermoFisher A1113803) starting at 24 hours post-transduction for at least 48 hours for HEK293FT cells and 5 days for THP1. For CaRPool-seq experiments (Figures 1 and 2) and direct capture Perturb-seq experiments (Figure 3), we transduced 3×10^6^ cells HEK293FT or NIH/3T3 cells. For screens conducted in THP1 cells (Supplementary Figures 6 and 7, and Figure 4) we transduced at least 12×10^6^ cells per condition. HEK293FT and NIH/3T3 cells were selected for at least two days, and THP1 cells for at least 5 days. For single-cell screens we selected conditions with < 10% survival 48 hours post puromycin selection or fraction of GFP-positive cells below 10% (MOI < 0.1), assuring high coverage (>1000x representation) and a single integration probability >95%. Pooled screens were conducted with MOIs between 0.13 and 0.45 always maintaining coverage of >1000x. Cas13d expression was induced using 1μg/ml doxycycline (Sigma D9891). Cells were maintained with doxycycline, blasticidin, and puromycin until the single-cell experiment. During this period, the cells were passaged every 2-3 days into fresh media supplemented with doxycycline, blasticidin, and puromycin.

### Flow cytometry

Puromycin-selected cells were harvested 2-7 days after selection start (HEK cells) or Cas13d induction (THP1 cells). (d)Cas9 sgRNAs were evaluated 7 days post-transduction. Cells were stained for the respective cell surface protein for 30 min at 4°C and measured by fluorescence-activated cell sorting (Sony SH800) (BioLegend: CD46 clone TRA-2-10 #352405 - 3μl per 1×10^6^ cells; CD55 clone JS11 #311311 - 5μl per 1×10^6^ cells; CD71 clone CYIG4 #334105 - 4μl per 1×10^6^ cells, CD11b clone ICRF44 #301322 – 2μl per 1×10^6^ cells). For flow cytometry analysis (FlowJo v10), cells were gated by forward and side scatters and signal intensity to remove potential multiplets. If present, cells were additionally gated with live-dead staining (LIVE/DEAD Fixable Violet Dead Cell Stain Kit, Thermo Fisher L34963). For each sample, we analyzed at least 5,000 cells. If cell numbers varied, we always subsampled (randomly) all samples to the same number of cells within an experiment.

### Bulk guide RNA detection

Puromycin-selected cells were harvested 48 hours or more after selection start. RNA was extracted from ~1×10^6^ cells (Zymo Direct-zol RNA microPrep). Total RNA was reverse transcribed using a capture sequence-specific reverse transcription primer along with an oligo(dT)V_30_ primer (400ng RNA, 4μl 5x RT Buffer, 1μl SuperasIN, 1μl dNTPs (10mM each), 1μl Maxima H Minus RT enzyme, 1.5μl 10μM Template Switch Oligo, 1μl 10μM oligo(dT)V_30_ primer, 1μl 10μM gRNA capture sequence primer, 20μl reaction volume; 53°C/90min, 70°C/15min). The Cas13 crRNA and *GAPDH* mRNA was amplified from 1:2 diluted cDNA using partial TSO and capture sequence primers or gene-specific primers (3μl cDNA, 10μl KAPA 2x master mix, 1μl primer (10 μM each), 5μl H_2_O; PCR conditions: 98°C/45s, 18x[98°C/20s, 60°C/10s, 72°C/10s], 72°C/5min). Oligonucleotides are provided in **Supplementary Table 2**.

### Cas13d gRNA off-target prediction

To identify potential Cas13d gRNA off-targets (alternative binding sites) we first aligned gRNAs to the human transcriptome (Grch38 cdna.all and ncRNA from emsembl release 97) using blastn (megablast) with the following parameters (-strand minus -max_target_seqs 10000 -evalue 10000 -word_size 5 - perc_identity 0.7). Secondly, candidates were further filtered to match with at least 17 bases, as shorter matches do not lead to target knockdown, and show a blastn e.value of < 100. In Supplementary Figure 3E, we demonstrate that despite the potential for reduced off-target binding, we do not observe transcriptomic perturbation for these genes.

### Bulk RNA-seq

To more carefully test for the potential of Cas13 to introduce off-target effects, we performed a bulk RNA-seq experiment, where we expect to have additional power to detect differential expression for lowly expressed transcripts. For the bulk RNA-seq experiment, we performed CD55 knockdown (Cas13d cells, KRAB-dCas9-MeCP2 cells) or knockout (Cas9-nuclease cells) using three individual targeting (s)gRMAs and three individual non-targeting (s)gRMAs. Monoclonal HEK293FT cell lines were transduced with a guide expressing lentivirus (MOI 0.2-0.5) in three independent transductions. Puromycin selection was started 24 hours post-transduction (1μg/mL), and Cas13d expression was induced (1μg/ml Doxycycline). Seven days post-transduction, we confirmed efficient CD55 targeting using flow cytometry, and total RNA was extracted (Zymo Direct-zol RNA microPrep). We performed a modified version of the Smart-seq2 protocol using 100ng purified total RNA input (https://www.protocols.io/view/barcoded-plate-based-single-cell-rna-seq-nkgdctw). Bulk RNA-seq samples were processed with Drop-seq tools v1.0 ^28^ using a hg19 reference. On average we obtained 352611 UMI (+/−72067 UMI) per sample. Differential gene expression was assessed with Seurat’s DESeq2 ^29^ implementation in FindMarkers.

### Pooled CRISPR screens in THP1 cells

Experimental procedures for performing multiplexed Cas13d screens in bulk were performed as described before ^11^, with minor modifications. THP1 cells were transduced and puromycin-selected as described above. Cas13d expression was induced after cells were fully selected (1μg/mL Doxycycline). Growth medium with fresh puromycin, blasticidin, and doxycycline was replenished every 2-4 days, and cells were split as needed always maintaining a guide representation of >1000x.

For the single guide RNA pooled screen, we collected a 1000x representation at 7 and 14 days post Cas13d induction and before sorting. After two weeks (14-16 days) we stained 15 million cells (~3000x representation), using FcX-blocking buffer (BioLegend #422302; 10 min at room temperature) and followed by either CD11b (BioLegend clone ICRF44 #301322 – 4μl per 1×10^6^ cells) or CD14 (BioLegend clone HCD14 #325608– 4μl per 1×10^6^ cells) staining (30 min at 4°C), and finally resuspending cells on PBS with DAPI (Sigma #D9542 - 0.4μg ml−1) to detect any apoptotic or dead cell. We sorted the cells (Sony SH800) based on their signal intensities (CD11b or CD14: lowest 10-15%, and highest 10-15%). Cells were PBS-washed and frozen at −80°C until sequencing library preparation. In total, we prepared four independent transductions (two MOIs and two alternative direct repeats), performed CD14 sorts for all four transduction replicates, and CD11b sorts for three transduction replicates collecting 1×10^6^ to 1.5×10^6^ cells per bin.

For the combinatorial targeting pooled screen, we prepared three transduction replicates (MOI 0.13-0.2). Eight days post Cas13d induction, we collected an input representation (>1000x coverage) and stained 20-30mio cells with FcX-blocking, CD11b, and DAPI as described above. We sorted the cells based on their CD11b signal intensity (lowest 15%, and highest 15%). Cells were PBS-washed and frozen at −80°C until sequencing library preparation. Library preparations for the single gRNA pooled screen were done as described before ^11^. For the combinatorial targeting pooled screen, we adopted a PCR strategy similar to the CaRPool-seq bcgRNA readout. Pooled screen readout PCR1 remained unchanged. In PCR2, we amplified the 15bp barcode sequence using a soluble Nextera-Read1-CS1 feature capture primer including an optional 28 randomized bases mirroring UMI and cell barcode, and RPIx Read2 i7 index primer. The amplicon was completed in PCR3 using Feature SI primer 2 (10x Genomics) and P7 primer.

### Pooled CRISPR screen analysis

Reads were demultiplexed based on Illumina i7 barcodes present in PCR2 reverse primers using bcl2fastq and, if applicable, by their custom in-read i5 barcode using a custom python script. For the single gRNA pooled screen, read1 sequencing reads were trimmed to the expected gRNA length by searching for known anchor sequences relative to the guide sequence using a custom python script. For the combinatorial pooled screen, we extracted the first 15 bases in read2. For the single gRNA pooled screen, we collapsed (FASTX-Toolkit) processed reads to count perfect duplicates followed by string-match intersection with the reference to retain only perfectly matching and unique alignments (average mapping rate 82.3%; median gRNA count 167). For the combinatorial pooled screen, pre-processed reads were either aligned to the barcode reference using bowtie ^30^ (v.1.1.2) with parameters -v 1 -m 1 --best –strata (average mapping rate 97%; median barcode read count 635; 1 barcode was not detected in input samples). For each dataset, raw counts were normalized using a median of ratios method as in DESeq2 ^29^ and batch-corrected using combat implemented in the SVA R package ^31^. Guide RNA and barcode enrichments were calculated building the count ratios between a sorting bin or timepoint and the indicated reference sample followed by log_2_-transformation (log_2_FC). For every single gRNA or bcgRNA, we considered the mean log_2_FC across replicates. For the single gRNA pooled screen, used the four best performing gRNAs per target gene to calculate the mean log_2_FC, where we determined best as either highest or lowest dependent on the sign of the mean enrichment across all ten gRNAs. As we have previously described ^11^, we noticed that log_2_FC enrichments were generally more pronounced in samples using the enhanced DR. Consistency between replicates and selected gRNAs was estimated using robust rank aggregation (RRA) ^32^. For the combinatorial pooled screen, we calculated the mean of both replicate arrays per gene pair. We noticed GFI1 g2 did not lead to strong effects in the pooled screen and in the CaRPool-seq experiment. The technical replicate arrays including GFI1 g2 were removed in all analyses. Enrichments are available in **Supplementary Tables 5** and **6**.

### Direct capture Perturb-seq

Monoclonal CRISPR-Cas effector protein-expressing cell lines (Cas9-nuclease, KRAB-dCas9, KRAB-dCas9-MeCP2) were infected with one of six sgRNA pools (KO-1, KO-2, KO-3, or CRISPRi-1, CRISPRi-2, CRISPRi-3) (**Supplementary Table 2**), providing 1-3 sgRNAs in a single vector and a total of nine cell line pool combinations. Cell survival after selection ranged between 1.7 and 5.5% (MOI < 0.1) assuring a high single integration probability. Viral titers were confirmed by measuring the fraction of BFP-positive cells for pools that have received vectors carrying 2+ sgRNAs using flow cytometry. Cells were passaged every 2-3 days continuously receiving puromycin and blasticidin at each split maintaining high sgRNA representation (>1000x coverage). We confirmed that >98% of cells were BFP-positive prior to the 10x experiment. We performed 10x V3.0 (Chromium Single Cell 3’ Gene Expression v3 with Feature Barcoding technology for CRISPR screening, #1000074, #1000075, #1000079) twelve days post-transduction. Cells were stained with a pool of five TotalSeq-A antibodies (0.75ug per antibody per 2×10^6^ cells) (**Supplementary Table 3**) following the CITE-seq protocol ^12^. In addition, we used Cell Hashing ^21^ (**Supplementary Table 3**) to track the nine cell line pool combinations. Before the run, cell viability was determined (≥ 96%). We ran one 10x lane, leveraging our hashed experimental design to overload with 38,600 cells. mRNA, sgRNA feature, hashtags (Hashtag-derived oligos, HTOs), protein (Antibody-derived oligos, ADTs) libraries were constructed by following 10x Genomics Cell-hashing and CITE-seq protocols ^12,21^. All libraries were sequenced together on one NextSeq 75 cycle high-output run.

### Direct capture Perturb-seq analysis

Sequencing reads coming from the mRNA library were mapped to the *hg38* (ensembl v97) genome reference using *Cellranger* (v3.1). Guide RNA reads were mapped simultaneously to a sgRNA feature reference (**Supplementary Table 1**). Prior to feature mapping, we performed 5’ adapter trimming using cutadapt to account for varying lengths of poly-G tracks five prime to the feature (first -g AAGCAGTGGTATCAACGCAGAGTACAT -O 5; then −O 1 -e 0 -g XGGGGGGGGGG) and trimmed the resulting sgRNA reads to a length of 18 bases. To generate count matrices for HTO and ADT libraries, the *CITE-seq-count* package (v1.4.2) was used (https://github.com/Hoohm/CITE-seq-Count). Count matrices were then used as input into the Seurat R package (v4.0) ^33^ to perform downstream analyses. We detected 16842 cells. HTO and sgRNA counts were normalized using the centered log-ratio transformation approach, with a margin = 2 (to normalize across cells instead of across features). To identity assign experimental conditions to cells and remove cell doublets, we used the *HTODemux* function in Seurat, with default parameters.

We customized *HTODemux* to return identities of second and third sgRNA assignments without changing the underlying modeling approach. We flagged cells with an incorrect number of expected sgRNAs numbers based on the HTO pool assignment. Furthermore, we flagged cells with an unexpected combination of sgRNAs not present in the sgRNA pool used to transduce the cells.

For the analysis shown in Figure 3, we only retained cells with the correct sgRNA numbers and identities. ADT counts were log-normalized, before running ScaleData with do.scale = FALSE and vars.to.regress = “PerturbSeq.approach”. PCA was performed on the log-normalized and regressed ADT counts using all five features, followed by UMAP dimensional reduction using four dimensions. To compare target knockdown across Perturb-seq approaches for NT-cell and cells that received all three (s)gRNAs (CD46, CD55, CD71), we normalized cellular ADT counts using median of ratios across ADT features that were not targeted (CD29, CD56) to derive a scaling factor per cell, and divided the normalized ADT counts by the mean ADT counts in NT cells for each Perturb-seq approach.

### Cas13 RNA Perturb-seq (CaRPool-seq) library preparation

We used Cas13 CRISPR array configurations of type X (extra PCR handle) as shown in **Figure 1a** and **Supplementary Figure 2**. Specifically, the bcgRNA was placed in the last array position and entailed a spacer sequence composed of a five-prime Illumina smallRNA PCR handle, a 15 base pair long barcode, and a three-prime capture sequence 1 (CS1) compatible with 10x Feature Barcoding technology. This composition allowed the specific amplification of a bcgRNA amplicon with a unique combination of forward and reverse primers. Moreover, usage of the Illumina 5’ PCR handle allows for efficient sequencing of the bcgRNA amplicon with the first base of Read 2 being the first barcode base. While this configuration has many advantages, other PCR primer sequences or capture sequences may be possible.

CaRPool-seq experiments were conducted using the 10x Genomics 3’ kit (Chromium Single Cell 3’ Gene Expression v3 with Feature Barcoding technology for CRISPR screening, #1000074, #1000075, #1000079). Library construction for bcgRNA derived oligos is outlined in **Supplementary Figure 2** and largely followed 10x Genomics user guide CG000184 Rev C for Feature Barcoding with some modifications. Specifically, we eluted the GEM-RT in 33ul and added 2μl containing 0.4μM ADT additive primer (for bcgRNAs and ADTs) and 0.2μM HTO additive primer prior to cDNA amplification. The cDNA was purified using 0.6x SPRI cleanup for mRNA fraction. The supernatant containing ADT, HTO, and bcgRNA cDNA was purified was purified by adding another 1.4x SPRI (0.6+1.4 = 2x SPRI) followed by an additional 2x SPRI clean up. The purified short fragments were split evenly into three pools (e.g. 3 x 20μl). One pool each was used for HTO and ADT library construction as described below. Half of the remaining pool (10μl) was used to construct the bcgRNA library using two PCR recipes. PCR1 adds Illumina P5 and P7 handles to the bcgRNA amplicon (100μl PCR1: 50μl 2x KAPA Hifi PCR Mastermix, up to 45μl bcgRNA PCR template, 2.5μl Feature SI Primers 2 10μM, 2.5μl TruSeq Small RNA RPIx primer (containing i7 index) 10μM; 95°C 3min, 12x[95°C 20sec, 60°C 8sec, 72°C 8sec], 72°C 1min). PCR2 amplifies with P5 and P7 primers (100μl PCR2: 50μl 2x KAPA Hifi PCR Mastermix, up to 45μl purified PCR1 product, 2.5μl P5 primer 10μM, 2.5μl P7 primer 10μM; 95°C 3min, 4x[95°C 20sec, 60°C 8sec, 72°C 8sec], 72°C 1min). The final bcgRNA amplicon has a length of 203bp and can be sequenced on a standard Illumina sequencing platform with standard Illumina sequencing primers (≥ 28 cycles Read1 and ≥15 cycles Read 2) (**Supplementary Figures 2** and **3a**).

### CaRPool-seq experiments

We transduced and treated Cas13d-NLS expressing HEK293FT, NIH/3T3, or THP1 cells as described above. In the species mixing, we used a pool of three bcgRNAs per species together with non-targeting gRNAs. The HEK293FT CaRPool-CITE-seq experiment included 29 CRISPR arrays barcoding a diverse set of array configurations around four gRNAs that allowed us to assess gRNA positioning within the CRISPR array, effects of the relative gRNA amount per cell, and combinatorial targeting of multiple RNA transcripts. All gRNA and bcgRNA sequences are provided in **Supplementary Table 1**. CaRPool-seq species mixing and CaRPool-CITE-seq were conducted simultaneously in one lane of 10x Genomics 3’ v3 kit. CaRPool-seq was performed on THP1 cells five days post Cas13d induction (1μg/mL Doxycycline) using four lanes of a10x Genomics 3’ v3 kit. THP1 CaRPool-seq library design and cloning were described above. Prior to the runs, cell viability was determined ≥ 95% for each experiment.

The HEK293FT CaRPool-seq experiment was stained with a pool of five TotalSeq-A antibodies (0.75ug per antibody per 2×10^6^ cells) (**Supplementary Table 3**) as following the CITE-seq protocol ^12^. Similarly, THP1 cells were first treated with FcX-blocking buffer (BioLegend #422302; 10 min at room temperature), before staining cells with a pool of 22 TotalSeq-A antibodies (**Supplementary Table 3**). To keep track of the experiment identity and identify multiplets, samples were hashed (subsequent to CITE-seq antibody staining) (**Supplementary Table 4**) following the Cell Hashing protocol ^21^. mRNA, hashtags (Hashtag-derived oligos, HTOs), protein (Antibody-derived oligos, ADTs) libraries were constructed by following 10x Genomics Cell-hashing and CITE-seq protocols ^12,21^.

Species mixing and HEK293FT CaRPool-seq experiment libraries were sequenced together on one NextSeq 75 cycle high-output run. THP1 CaRPool-seq libraries were sequenced on NovaSeq6000 using the XP S4 2×100 v1.5 workflow. Sequencing reads coming from the mRNA library were mapped to a joined genome reference of *hg38* (ensemble v97) and *mm10* using the *Cellranger* Software (v3.0.1), or to *hg38* using *Cellranger* v6.0.0 for the THP1 experiment. Barcode guide RNA library reads were mapped simultaneously to a barcode reference (**Supplementary Table 1**) using *Cellranger*. To generate count matrices for HTO and ADT libraries, the *CITE-seq-count* package (v1.4.2) was used (https://github.com/Hoohm/CITE-seq-Count). Count matrices were then used as input into the *Seurat* R package (v4.0) ^33^ to perform all downstream analyses.

### CaRPool-seq data analysis

Cells from species-mixing and HEK293FT CaRPool-seq experiments were processed together. Cells with <2,500 UMI were removed. HTO and bcgRNA counts were normalized using the centered log-ratio transformation approach, with a margin = 2 (to normalize across cells instead of across features). To identity cell doublets and assign experimental conditions to cells, we used the *HTODemux* function. Only human cells were hashed, with mouse NIH/3T3 cells being the only cell population without a hashtag. We removed all hashing doublets within the CaRpool-CITE-seq experiment (HTO-01 to HTO-08) and to human cells in the species mixing experiment (HTO-10). In addition, we removed all cells labeled with a single HTO-01 to HTO-08 if the fraction of mouse reads was >10%, and cells without any HTO if not at least 10% mouse reads were present. Like this, we removed all doublets between CaRPool-seq species mixing and CaRPool-CITE-seq experiments while retaining potential collisions/doublets between mouse and human cells as part of the CaRPool-seq species mixing branch. At this point, the experiment was split into two separate objects. For the CaRPool-seq species mixing experiment, we determined species identity by quantifying the fraction of human reads for RNA and for the species-specific bcgRNAs (human: > 0.9, mouse: < 0.1, collision: 0.9 to 0.1). For the HEK293FT CaRPool-CITE-seq experiment RNA counts were log-normalized using the standard Seurat workflow after removing all mouse features and RNA counts. Barcode guide RNA identity was determined using *MultiSeqDemux(autoThresh = T)*. Cells without bcgRNA assigned and cells with multiple bcgRNA assignments were removed.

For the THP1 experiment we detected 52,496 single cells (nFeature_RNA > 1000, nFeature_RNA < 8000, percent.mt < 20) after HTO demultiplexing using *HTOdemux* as described above. Model-based bcgRNA assignments (*HTODemux* or *MultiSeqDemux*) did not yield satisfying results supported by the observed phenotypic changes, likely due to model limitations imposed by the high number of bcgRNA features. Instead, we assigned bcgRNAs to single cells by applying the following rules:

We compared UMI counts for the bcgRNA with the highest UMI count (g1) to, if present, the second detected bcgRNA (g2). bcgRNA counts for g2 may derive from spurious counts arising from library preparation, or from integration of more than one viral element (bcgRNA multiplet). We considered cells with g1 < 5 as Negative. We assigned g1 if: 1) g1 = {5-9} and g2 = {0-1}, or 2) g1 > 9 and g1/(g1+g2) >0.8 and g2 < 11. All other cells were considered bcgRNA multiplets. We assigned 31,308 with a single bcgRNA. Comparing differential gene expression results for technical replicates embedded in the CaRPool-seq library, we noticed GFI1 g2 did not lead to upregulation of CD11b ADT and did not lead to upregulation of the expected gene expression signature. We removed all cells with GFI1 g2 (n=601). Changes in cell surface protein ADT levels for gene pair or individual CRISPR array were calculated using Wilcoxon’s rank-sum test in FindMarkers relative to NT control cells. Changes were determined by repeating the differential expression analysis ten times with <= 30 randomly samples cells per cell group to account for differing numbers of cells.

### Extrapolation of sgRNA detection in direct capture Perturb-seq experiments

We used published sgRNA assignment rates for single and dual sgRNA targeting using direct sgRNA capture via Feature Barcoding technology ^6^. We determined the mean sgRNA assignment rate to be 80.9% by averaging the assignment rate for exactly one sgRNA (80% in single guide experiments) and taking the square root of the assignment rate of exactly two sgRNAs per cell (67% in dual guide experiments). In our simulation, we assume that a single viral particle will be taken up by a cell during a low-MOI infection. A single integration event may deliver up to three sgRNAs that are independently expressed, similar to the two sgRNA experiments described previously ^6^. We assume that sgRNA-detection for each sgRNA is an independent event. These can be modeled by multiplying detection and editing probabilities *p* by the number of sgRNA feature *n* (*p^n^*). The resulting curve shows the fraction of cells that have received exactly *n* sgRNAs (grey line in Figure 3b).

### Modeling of genetic interactions (GI) in single-cell data

To decompose transcriptomic profiles of double perturbation, we used a linear regression model as previously introduced ^7^ and implemented it in R. First, we z-scaled the log-normalized gene expression counts for all cells with respect to the mean and standard deviation of the control group (non-targeting cells). In this way, we have subtracted the baseline expression profiles from each cell and can directly compare the deviation from each perturbation to NT conditions. Next, we grouped cells by gene pair and calculated pseudo-bulk z-scaled profiles [single perturbations (a, b), and double perturbation (ab)] by calculating the mean across cells for each feature. The average NT-cell profile returns a vector of all zeros. We generated average profiles for 1,530 genes with an average UMI count > 0.5. We included gene pairs when all cell groups were represented by at least 25 cells (Examples in Figure 4e-i indicate cell count in parathesis).

As previously introduced ^7^, we model the average z-scale profiles using:

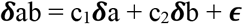

With ***δ*** a is the pseudobulk z-scaled profile for cells assigned to single perturbation a, ***δ*** b is the pseudobulk z-scaled profile for cells assigned to single perturbation b, and ***δ*** ab is the pseudobulk z-scaled profile for cells assigned to double perturbation ab. c_1_ and c_2_ are constants fitted to the data indicating the relative weight of ***δ*** a and ***δ*** b profiles. The vector ***ϵ*** collects the residuals to the model fit. In our plots, a is the first gene in the gene pair, and b is the second gene. c_1_ corresponds to a, and c_2_ to b.

We implemented the previously-introduced model-fitting procedure ^7^, using the *rlm* function from the MASS package, and extracted the mean coefficients (c_1_ and c_2_) and residual error ***ϵ***. We collected six measures to evaluate the fit as described before ^7^ (*dcor* function from energy package):

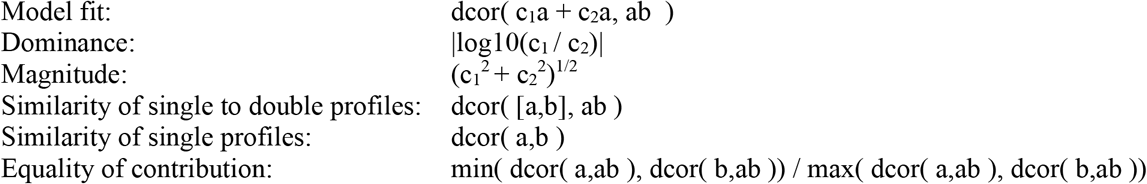

Each feature and its interpretation are described in detail here ^7^. Features were scaled (margin = 2) prior to hierarchal clustering (dist = euclidean, methods = ward) to generate a dendrogram, shown in Figure 4D. For clarity, the heatmap in Figure 4d shows unscaled values.

The example interactions shown in Figure 4e-i depict the union of top 20 differentially expressed genes for each cell group (a, b, ab) relative to NT-cells (selected by p-value) derived using the Wilcoxon’s rank-sum test in FindMarkers. Model prediction and residuals are derived from the modeling approach described above. The color scale represents the average z-score normalized expression per gene pair.

## SUPPLEMENTARY TABLES

Supplementary Table 1: CaRPool-seq and Perturb-seq libraries

Supplementary Table 2: Oligonucleotides for molecular cloning

Supplementary Table 3: ADTs used in this study

Supplementary Table 4: HTOs used in this study

Supplementary Table 5: CD11b and CD14b pooled screen results

Supplementary Table 6: CD11b combinatorial pooled screen results

