## Supplementary Figures for "Efficient combinatorial targeting of RNA transcripts in single cells with Cas13 RNA Perturb-seq"

### SUPPLEMENTARY FIGURE 1

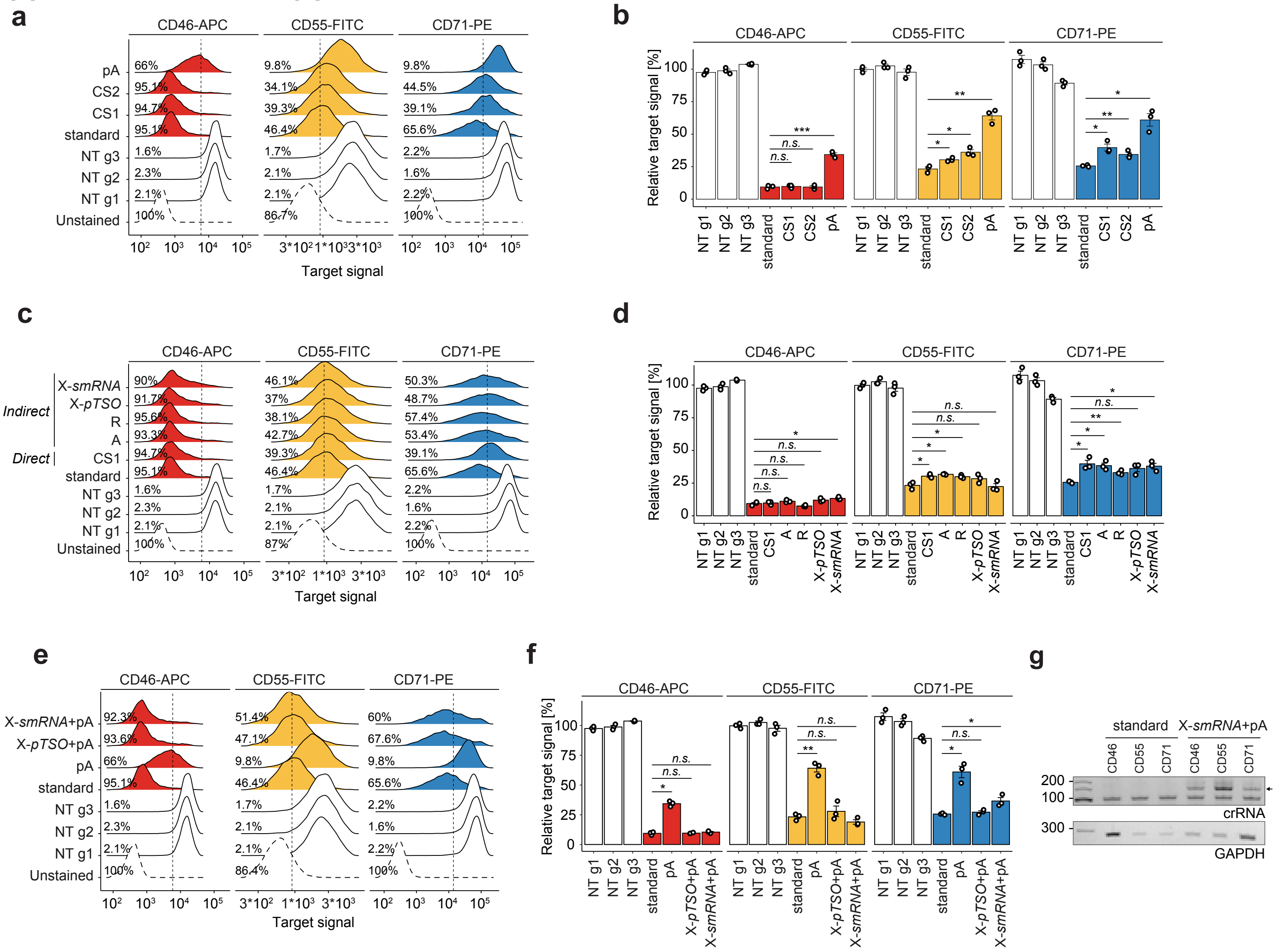


**Supplementary Figure 1: Direct guide RNA capture by addition of a 3’ common sequence to the Cas13d spacer RNA**

a) Density plots showing the CD46-APC, CD55-FITC and CD71-PE flow cytometry signal upon Cas13d-mediated knockdown with either regular gRNAs or a direct capture gRNA with one of three reverse transcription handles ($pA_{(30)}$ = polyA-tail of length 30, CS1 = 10x Genomics Capture Sequence 1, CS2 = 10x Genomics Capture Sequence 2, NT = non-targeting). Vertical lines mark the threshold for CD-protein negative cells ($2^{nd}$ percentile of NT cell populations), indicating the percent negative cells for one replicate experiment. Importantly, the Cas13-mediated function shows a unimodal response, suggesting limited cell-to-cell differences in target gene knockdown.

b) Summary analysis of three replicate experiments as shown in (a). Y-axis shows the mean fluorescent intensity (MFI) relative to the average of all NT cell populations. Direct capture constructs with CS1 or CS2 enable strong knockdown for CD46, but reduced knockdown for CD55 and CD71. Direct capture with pA-handle shows strongly reduced knockdown efficiency compared to regular gRNAs (standard condition). Two-sided t-test with * $p < 0.05$, ** $p < 0.01$, and *** $p < 0.001$.

c) Density plots showing the CD46-APC, CD55-FITC, and CD71-PE signal upon Cas13d-mediated knockdown with either regular gRNAs, a direct capture gRNA, or indirect capture construct of types A, R, and X as shown in Figure 1a. CS1 was used in all constructs with RT-handle. Type X was used with either a partial TSO (pTSO) PCR priming site or an Illumina smallRNA PCR-handle sequence. Vertical lines mark the threshold for CD-protein negative cells, indicating the percent negative cells for one replicate experiment.

d) Summary analysis of three replicate experiments as shown in (c). Y-axis shows the mean fluorescent intensity (MFI) relative to the average of all NT cell populations. Indirect capture constructs show strong target gene knockdown similar to regular gRNAs (standard condition) for all three target genes. The slight reduction in targeting efficiency in indirect guide capture may be explained by CRISPR array processing constraints. Two-sided t-test with * $p < 0.05$, ** $p < 0.01$, and *** $p < 0.001$.

e) Density plots showing the CD46-APC, CD55-FITC, and CD71-PE signal upon Cas13d-mediated knockdown with either regular gRNAs, a direct capture gRNA, or indirect capture construct of type X. Here, comparing the effect and placement of a polyA-tail RT-handle. Type X was used with either a pTSO or smallRNA PCR-handle sequence. Vertical lines mark the threshold for CD-protein negative cells, indicating the percent negative cells for one replicate experiment.

f) Summary analysis of three replicate experiments as shown in (e). Y-axis shows the mean fluorescent intensity (MFI) relative to the average of all NT cell populations. Indirect capture constructs show strong target gene knockdown like regular gRNAs (standard condition) for all three target genes. Target knockdown with direct capture through a polyA-tail sequence is limited. Two-sided t-test with * $p < 0.05$, ** $p < 0.01$, and *** $p < 0.001$.

g) PCR amplicons of reverse-transcribed crRNAs from lentivirally infected cells used in e. Indirect capture of Type-X crRNAs with smallRNA PCR-handle and polyA-tail (arrow) allowed for reverse transcription and amplification. These results show that indirect gRNA capture can be facilitated with polyA-tail capture as an alternative to CS1-based capture. X-type arrays are independent of template switching success.

### SUPPLEMENTARY FIGURE 2

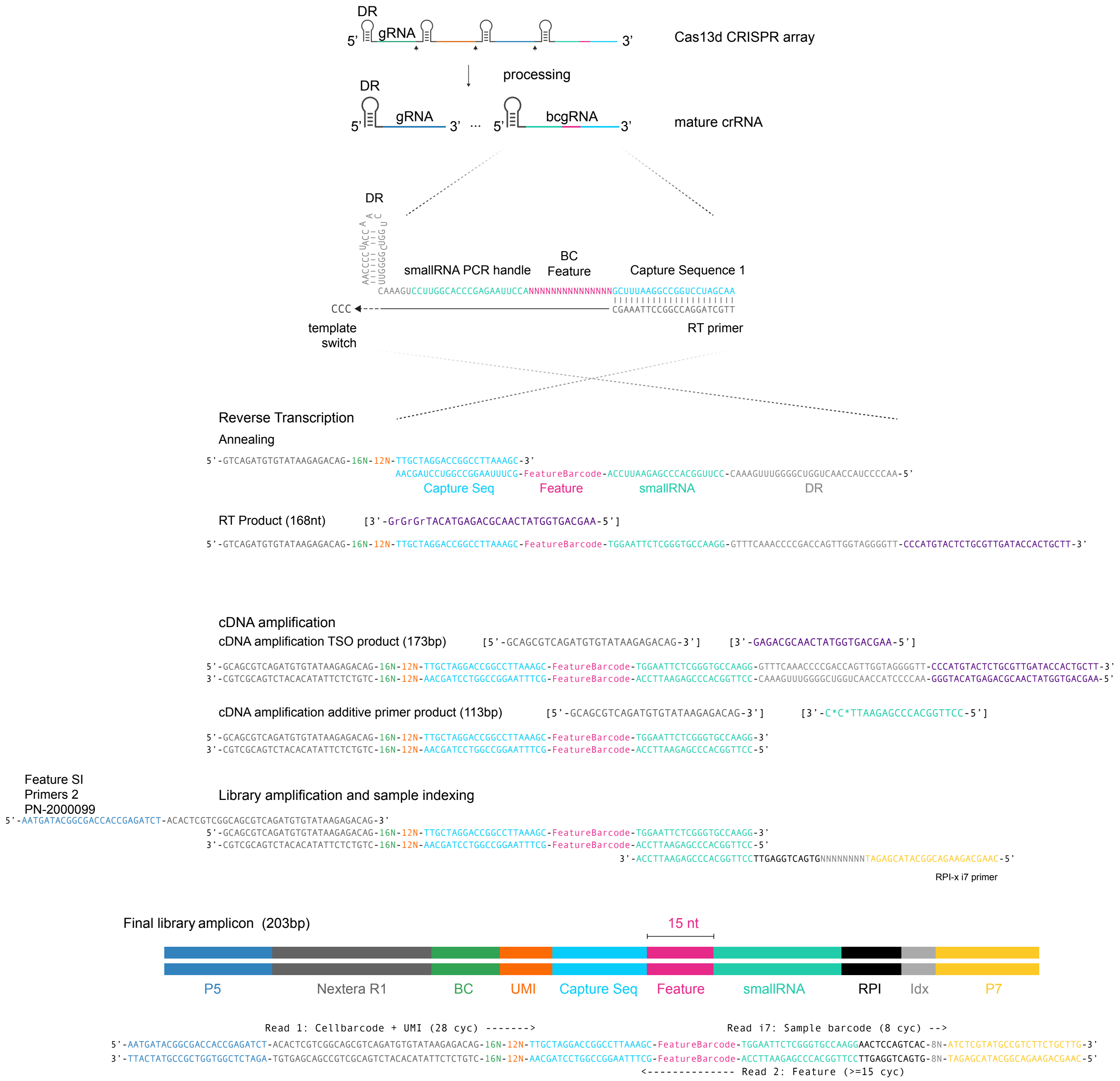


**Supplementary Figure 2: bcgRNA capture scheme adapted from 10x Genomics Feature Barcoding technology.**

### SUPPLEMENTARY FIGURE 3

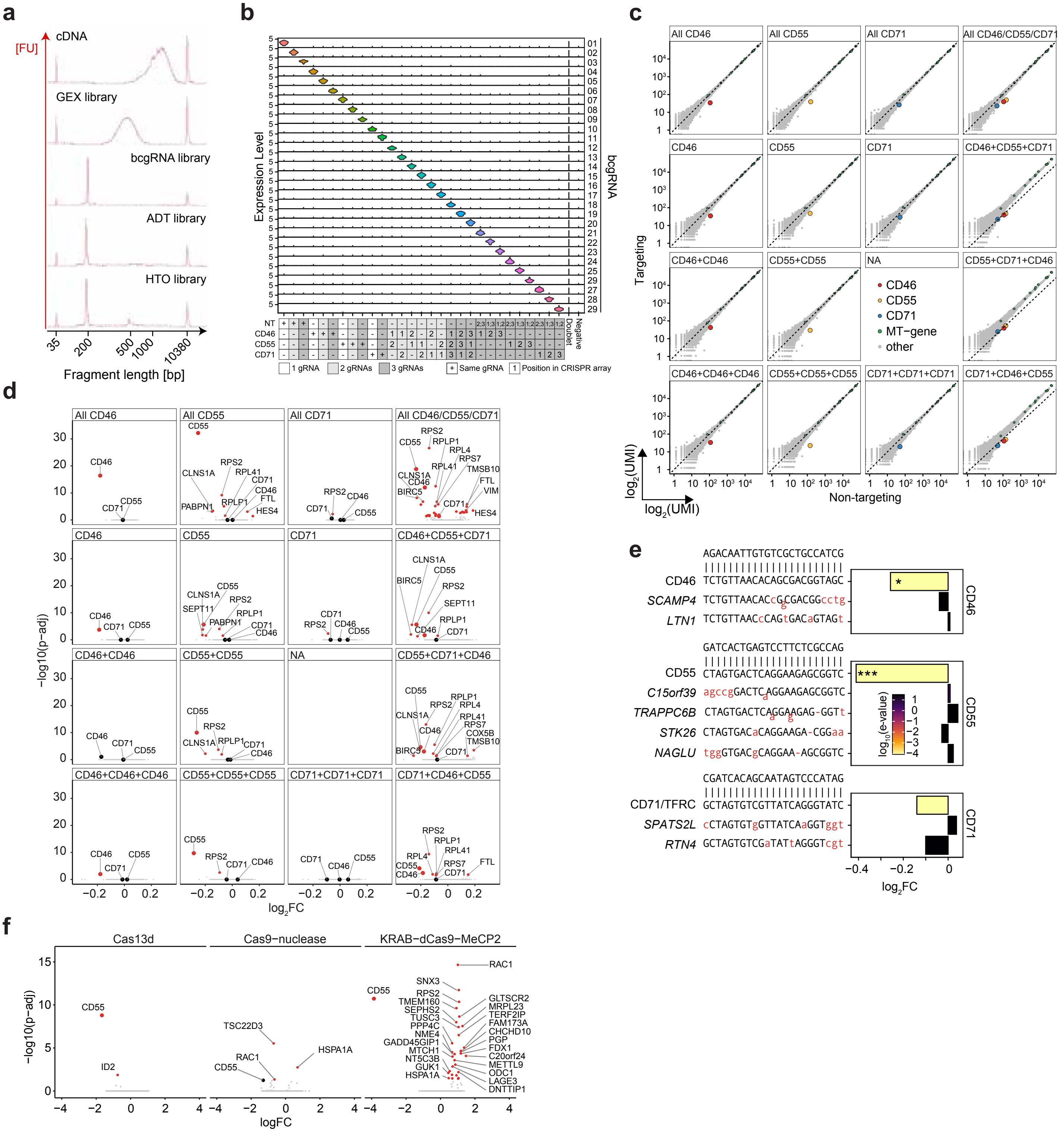

**Supplementary Figure 3: CaRPool-seq enables efficient bcgRNA capture and specific target RNA knockdown.**
a) Representative BioAnalyzer traces of cDNA and four jointly assayed modalities (GEX = gene expression, bcgRNA = barcode guide RNA, ADT = antibody derived tags, HTO = hashtag oligonucleotides). b) Stacked violin plot showing normalized bcgRNA UMI counts for cells grouped by assigned CRISPR array [total cells n = 9,355, cells with single bcgRNA n = 6,986, (74.7%)]. c) Scatterplots showing normalized pseudobulk RNA UMI count profiles of cells grouped by indicated CRISPR arrays (y-axis) and control NT-cells (x-axis). Respective target genes (CD46, CD55, CD71) are highlighted in color. Genes on the MT chromosome are colored green. CD71+CD71 was not included in the experiment. d) Volcano plots showing differential gene expression results cells grouped by indicated CRISPR arrays and control NT cells. Cells grouping is the same as in Figure S3c. The x-axis indicates log-transformed fold changes. The y-axis depicts -$\log_{10}$-transformed adjusted p-values (Wilcoxon test). Significantly differentially regulated genes (adjusted p-value < 0.05) are highlighted in red. e) Sites, and relative expression levels of gRNA-dependent predicted off-target transcripts from gRNAs targeting CD46, CD55 and CD71. Red letters indicate mismatches to cognate perfect match target site. E-values derived from Blastn. (Wilcoxon test * p.adj. < 0.05, ** p.adj. < 0.01, *** p.adj. < 0.001). f) Bulk RNA-seq result for Cas13d, Cas9-nuclease, and KRAB-dCas9-MeCP2 based targeting of CD55 using three independent CD55-targeting and NT (s)gRNAs, respectively. Volcano plots show differential gene expression results of CD55 targeting conditions relative to corresponding NT conditions grouped by indicated CRISPR effector protein. The x-axis indicates log-transformed fold changes. The y-axis depicts -log10-transformed adjusted p-values (DESeq2). Significantly differentially regulated genes (adjusted p-value < 0.05) are highlighted in red. The three approaches show a varying number of differentially expressed genes in addition to CD55 reduction (n=1 Cas13d, n=3 Cas9, n=30 KRAB-dCas9-MeCP2). Cas13d gRNA and Cas9 sgRNA efficiency is shown in *Supplementary Figure 4a* and *Supplementary Figure 5a*.

### SUPPLEMENTARY FIGURE 4

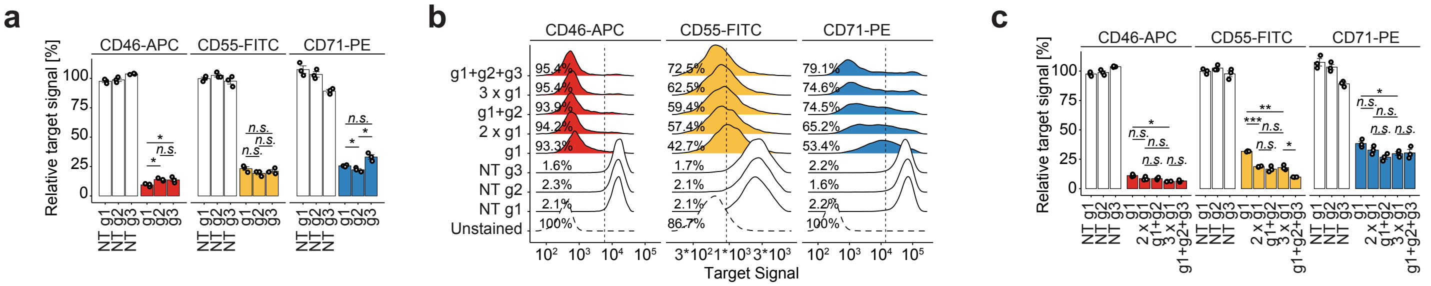


**Supplementary Figure 4: CaRPool-seq detects target RNA and protein knockdown for single guide RNAs**

a) Bar plots depicting CD46-APC, CD55-FITC, and CD71-PE signal upon Cas13d-mediated knockdown with three alternative gRNAs per target gene relative to the mean of three NT controls measured by flow cytometry. Y-axis shows the mean fluorescent intensity (MFI) relative to the average of all NT cell populations. Two-sided t-test with * $p < 0.05$, ** $p < 0.01$, and *** $p < 0.001$. Guide RNA g1 was used in CaRPool-seq experiments. Guide RNAs g2 and g3 are used in figures b and c.

b) Density plots showing the CD46-APC, CD55-FITC, and CD71-PE signal upon Cas13d-mediated knockdown with either 1, 2, or 3 copies of the same gRNA (g1) per CRISPR array or 2 and 3 alternative gRNAs (g2, g3). Vertical lines mark the threshold ($2^{nd}$ percentile of combined NT conditions) for CD-protein negative cells, indicating the percent negative cells for one replicate experiment. Shown is one representative replicate

c) Summary analysis of three replicate experiments as shown in c. Y-axis shows the mean fluorescent intensity (MFI) relative to the average of all NT cell populations. The Analysis suggests that target gene knockdown differences between the number of gRNAs per array are more pronounced than differences between gRNA identities with the same total count, given that gRNA efficiencies are comparable as shown in b. CRISPR arrays encoding multiple gRNAs against the same target may be used to further enhance target knockdown. Two-sided t-test with * $p < 0.05$, ** $p < 0.01$, and *** $p < 0.001$.

### SUPPLEMENTARY FIGURE 5

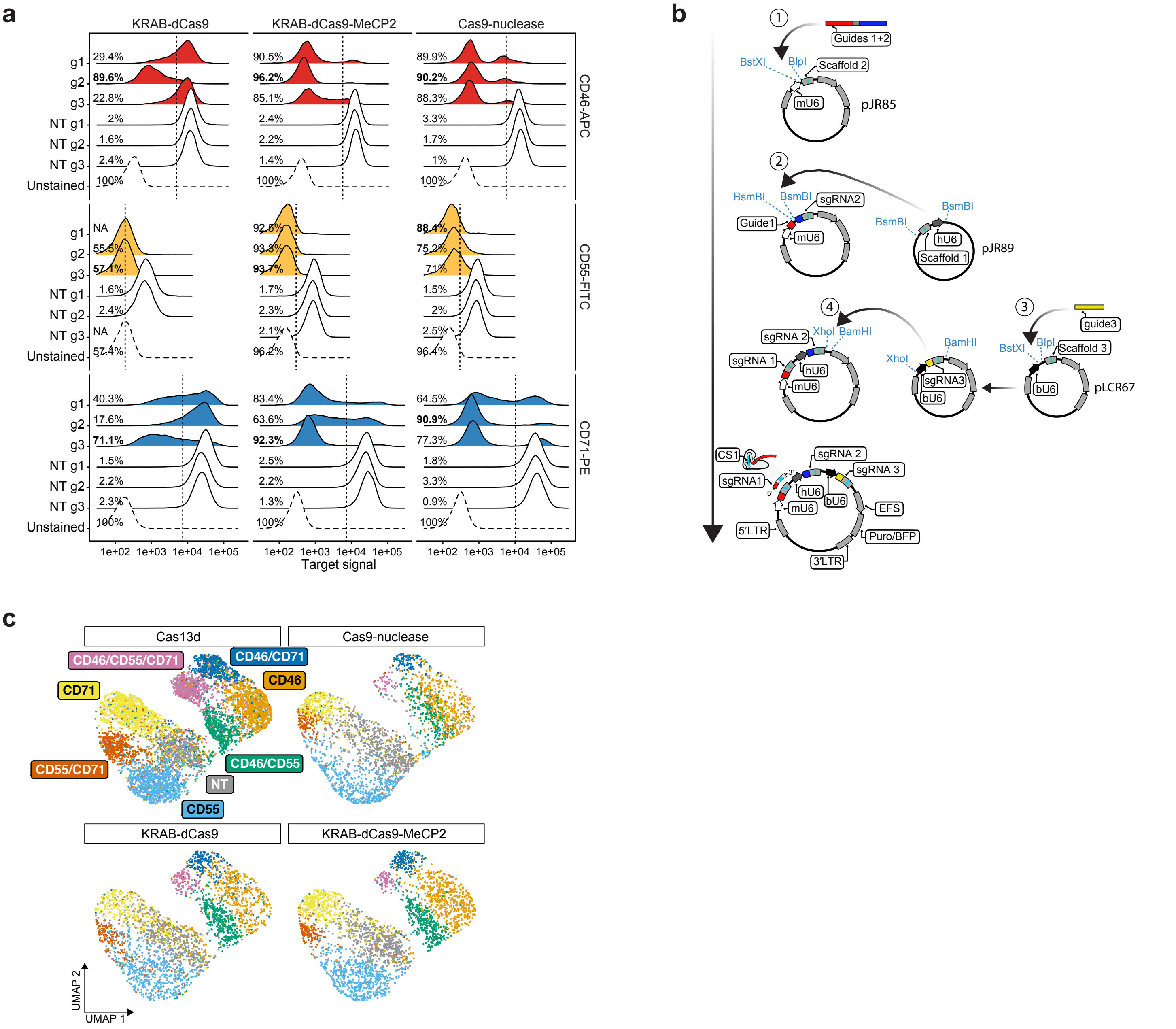


**Supplementary Figure 5: CaRPool-seq enables efficient combinatorial target RNA perturbation.**
a) Density plots showing the CD46-APC, CD55-FITC, and CD71-PE flow cytometry signal upon Cas9-nuclease mediated knockout (KO) and CRISPRi-mediated (KRAB-dCas9, KRAB-dCas9-MeCP2) knockdown with three alternative sgRNAs from established genome-wide KO [19] and CRISPRi [20] libraries. Vertical lines mark the threshold (2$^{nd}$ percentile of combined NT conditions) for CD-protein negative cells, indicating the percent negative cells for one replicate experiment. Single guide RNAs with the highest percentage of negative cells (*bold*) were selected for direct capture Perturb-seq experiments (NA = sgRNA not assayed).
b) Cloning strategy for triple sgRNA plasmid vectors. Dual sgRNA constructs were cloned as described before [6]. The third sgRNA was cloned behind a bovine U6 promoter using an alternative sgRNA scaffold tested before [6].
c) Protein level ADT-based clustering of single-cell expression profiles of merged CaRPool-CITE-seq (n = 6,986 cells) and Perturb-seq experiments using Cas9-nuclease (n = 2,836), KRAB-dCas9 (n = 2,911) or KRAB-dCas9-MeCP2 (n = 3,038) effector proteins as in Figure 3e. Cells are labelled by the assigned target gene combination based on detected bcgRNA or sgRNAs and split by Perturb-seq.

### SUPPLEMENTARY FIGURE 6

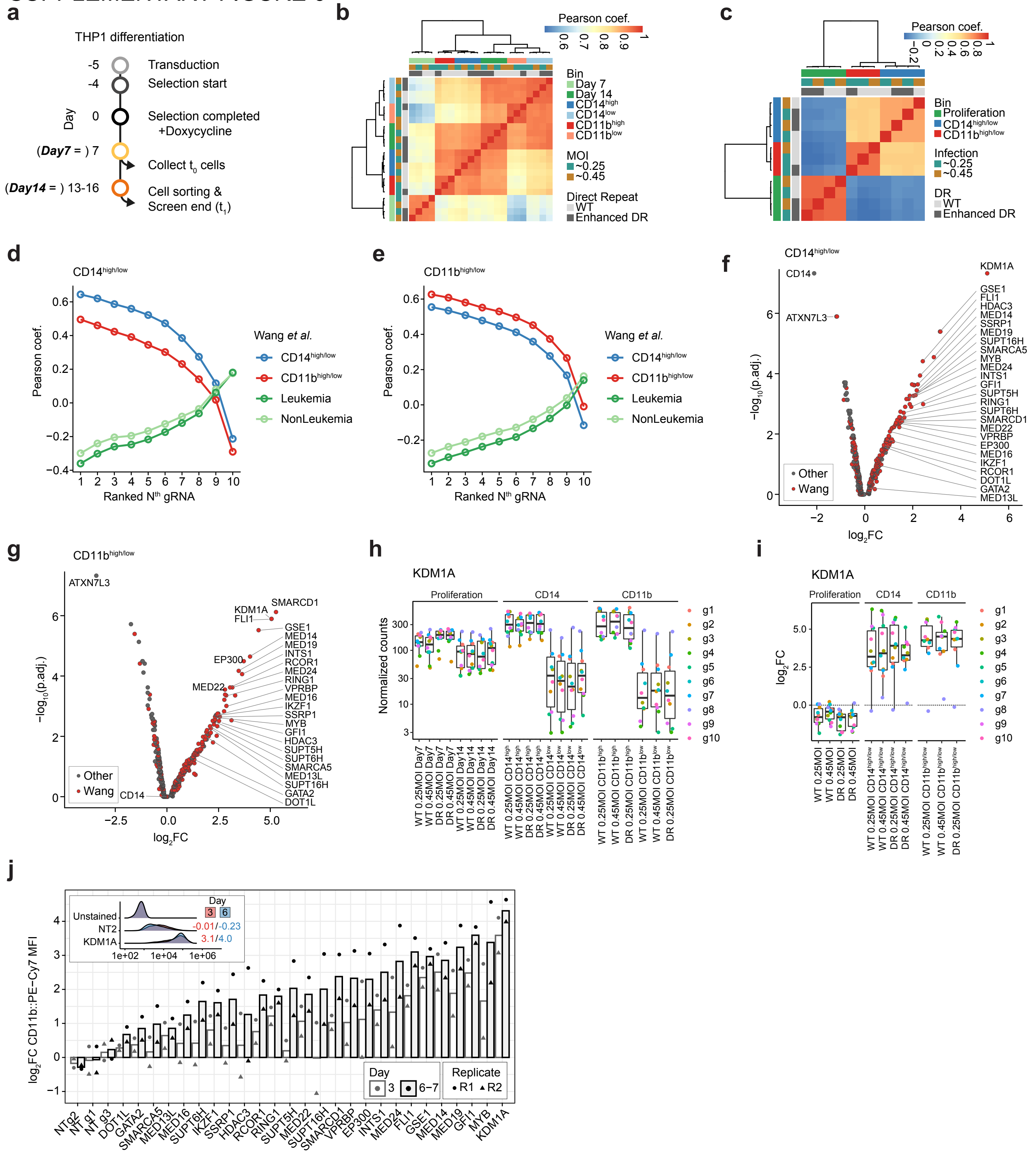

**Supplementary Figure 6: A pooled Cas13 screen identifies regulators of AML differentiation.**
a) Timeline for THP1 cell infections and pooled screen readouts for CD14 and CD11b fluorescent activated cell sorting (FACS). We transduced a pooled lentivirus library with 4800 gRNAs targeting 439 genes (10 gRNAs per gene) and 410 NT control gRNAs. Each gRNA was tested using two alternative Cas13d direct repeat (DR) sequences [wildtype (WT) or enhanced DR; see methods]. Cells were infected at two different MOIs for each DR. Cells were collected at Day seven (timepoint 0; $t_0$) and approximately Day 14 (range between day 14 and day 16; $t_1$) post Cas13d induction. On day 14, cells were sorted based on their cell surface protein expression into CD14 and CD11b high (top 15%) and low (bottom 15%) bins. At each time point, we collected and sorted cell populations with >1000x coverage. With alternative MOI infections and DR sequences used, we conducted four and three replicated phenotypic cell sorts for CD14 and CD11b.
b) Pearson correlation of normalized and batch corrected gRNA counts for all samples.
c) Pearson correlation of $log_2$-transformed gRNA enrichments of the population of interest relative to the corresponding control population ($CD11b^{high/low}$: gRNA counts in $CD11b^{high}$ bin divided by $CD11b^{low}$ bin; $CD14^{high/low}$: gRNA counts in $CD14^{high}$ bin divided by $CD14^{low}$ bin; Proliferation: gRNA counts at $t_1$ divided by $t_0$).
d) Correlation of $log_2$FC gene enrichments for CD14 upregulation ($CD14^{high/low}$) to enrichments presented in Wang et al.. Correlation analysis was repeated for every single ranked (see methods) gRNA, always considering only one ranked gRNA per target gene. This analysis indicated that, as expected, target gene enrichments for CD14 upregulation correlated best with CD14 target gene enrichments in Wang et al.. And that correlations were similarly high for the top-ranked 4-5 gRNA.
e) Similar analysis as presented in d showing result for target genes regulating CD11b enrichments.
f) Volcano plot showing gene enrichments for target genes regulating CD14 upregulation ($CD14^{high/low}$). Each gene is represented as the mean of the four top-ranked gRNAs across all replicate experiments (see methods). Y-axis shows -log10 transformed adjusted p-value derived from robust ranked analysis (RRA). Target genes selected based on previous results (Wang et al.) are shown in red. The 28 genes used in the subsequent CaRPool-seq experiment are highlighted. As expected, CD14 was the most depleted gene.
g) Similar analysis as presented in f showing results for target genes that lead to CD11b enrichment. CD11b-targeting gRNAs were not included in the gRNA library.
h) Example of gRNA counts across all collected samples for the KDM1A target gene.
i) Example of gRNA enrichments of counts shown in h across all comparisons according to comparisons described in c for the KDM1A target gene. The gRNA color legend applies to h and i.
j) Individual gRNA confirmations for 26 hit genes and three NT controls. Cas13d expressing THP1 cells were transduced with individual gRNAs expressing lentivirus targeting one out of 26 selected target genes. Three and six-to-seven days (technical replicates) cells were stained for CD11b and CD11b levels were recorded. Y-axis shows the CD11b::PE-Cy7 MFI relative to the average of all three NT control samples for the respective time point. Bars show the mean of two independent replicates. The inset shows CD11b::PE-Cy7 levels in KDM1A and NT targeted cells three and six days after Cas13d induction.

### SUPPLEMENTARY FIGURE 7

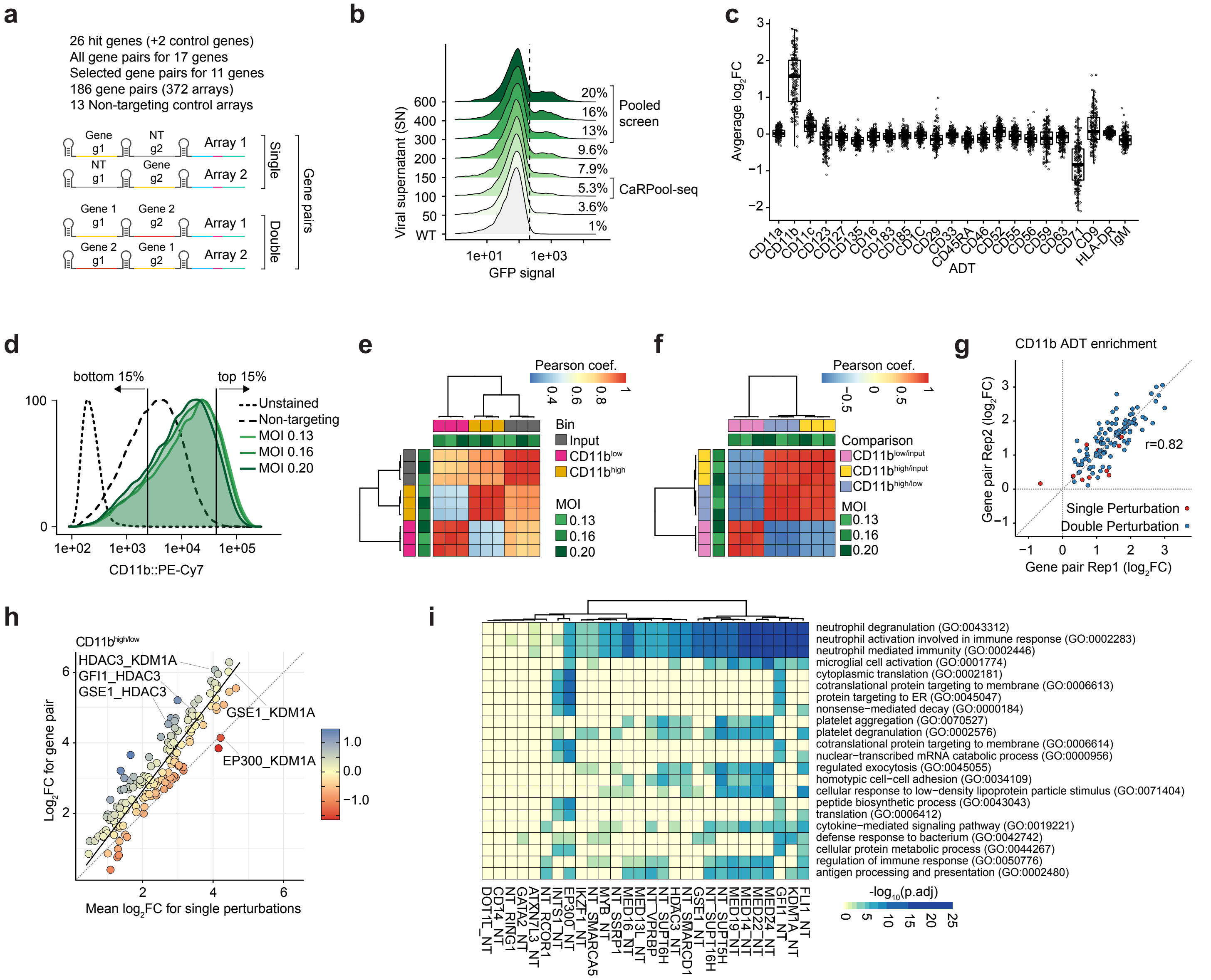


**Supplementary Figure 7: Combinatorial targeting of AML differentiation regulators.**

a) Library composition of CaRPool-seq and pooled screen for CD11b fluorescent activated cell sorting (FACS). We generated a pooled library with 385 bcgRNA CRISPR arrays, containing 158 gene pair perturbations (gene 1 – gene 2), 28 single-gene perturbations (gene - NT). Each single and dual perturbation is targeted by two sets of non-overlapping gRNAs (=technical replicate) and 13 non-targeting controls.

b) Titration of CaRPool-seq lentivirus four days after transduction. Viral constructs contain a Puromycin-2A-GFP expression cassette. Density plots show the percent GFP positive cells compared to the 99th percentile of WT control cells. The percent GFP-positive cells indicate the MOI. Low-MOI cells were used in the CaRPool-seq experiment, and high-MOI conditions were used for pooled screen readout.

c) Differential expression analysis for ADTs of cell surface proteins all assayed in the CaRPool-seq experiment. Each point represents results for one gene pair summarizing the effect across both independent CRISPR array replicates per gene pair (n=186, single and double perturbations). To account for differences in cell number per gene pair we calculated log2FCs as the mean of 10 samples of a random cell using 30 cells (or all cells for gene pairs < 30 cells) per gene pair relative to the same number of NT control cells. All but two gene pairs show elevated levels of CD11b.

d) Density plots showing the CD11b::PE-Cy7 signal of THP1 cells transduced with NT gRNAs or the CaRPool-seq library 8 days post Cas13d induction. We collected unsorted cells alongside sorting cells based on CD11b signal collecting the 15% of lowest and highest CD11b signal. We collected and sorted cell populations with >1000x coverage.

e) Pearson correlation of normalized and batch corrected bcgRNA counts for all samples.

f) Pearson correlation of $\log_2$-transformed bcgRNA enrichments of population of interest relative to the corresponding control population (CD11b$^{high/low}$: bcgRNA counts in CD11b$^{high}$ bin divided by CD11b$^{low}$ bin; CD11b$^{high/input}$: bcgRNA counts in CD11b$^{high}$ bin divided by unsorted input representation; CD11b$^{low/input}$: bcgRNA counts in CD11b$^{low}$ bin divided by unsorted input representation).

g) Comparison of $\log_2$FC CD11b ADT enrichments relative to NT cells in CaRPool-seq data comparing the two technical replicate gene pair arrays. Shown are all technical replicates where both arrays were represented by at least 25 cells (n=122 gene pairs).

h) Correlation of CD11b$^{high/low}$ $\log_2$FC enrichments of in dual perturbation cells and the mean $\log_2$FC of both single perturbation cells corresponding to the dual perturbation (n=158 gene pairs). Residuals indicate the distance to the average linear relationship. For each gene pair, we used the mean of both replicate CRISPR arrays.

i) EnrichR gene ontology analysis for biological processes (GOBP) for all single-gene perturbations (n=28) using up to 100 significantly (p<0.01) upregulated genes per gene pair compared to NT condition. Shown are the -$\log_{10}$-transformed adjusted p-values for GO-terms with p<0.00001 in at least one condition.
